# The stability of the primed pool of synaptic vesicles and the clamping of spontaneous neurotransmitter release relies on the integrity of the C-terminal half of the SNARE domain of Syntaxin-1A

**DOI:** 10.1101/2023.07.06.547909

**Authors:** Andrea Salazar-Lázaro, Thorsten Trimbuch, Gülçin Vardar, Christian Rosenmund

**Affiliations:** Department of Neurophysiology, Charité-Universitätsmedizin Berlin, Humboldt-Universität zu Berlin, and Berlin Institute of Health, 10117 Berlin, Germany; NeuroCure Excellence Cluster, Berlin, Germany

**Keywords:** SNARE domain, C-terminus half of the SNARE domain, Syntaxin-1(STX1)

## Abstract

The SNARE proteins are central in membrane fusion and, at the synapse, neurotransmitter release. However, their involvement in the dual regulation of the synchronous release while maintaining a pool of readily releasable vesicles remains unclear. Using a chimeric approach, we performed a systematic analysis of the SNARE domain of STX1A by exchanging the whole SNARE domain or its N- or C-terminus subdomains with those of STX2. We expressed these chimeric constructs in STX1-null hippocampal mouse neurons. Exchanging the C-terminal half of STX1’s SNARE domain with that of STX2 resulted in a reduced RRP accompanied by an increased release rate, while inserting the C-terminal half of STX1’s SNARE domain into STX2 lead to an enhanced priming and decreased release rate. Additionally, we found that the mechanisms for clamping spontaneous, but not for Ca^2+^-evoked release, are particularly susceptible to changes in specific residues on the outer surface of the C-terminus of the SNARE domain of STX1A. Particularly, mutations of D231 and R232 affected the fusogenicity of the vesicles. We propose that the C-terminal half of the SNARE domain of STX1A plays a crucial role in the stabilization of the RRP as well as in the clamping of spontaneous synaptic vesicle fusion through the regulation of the energetic landscape for fusion, while it also plays a covert role in the speed and efficacy of Ca^2+^-evoked release.

## Introduction

The molecular release machinery underlying synaptic vesicle fusion in mammalian central synapses consists of the SNARE complex at its core and several accessory proteins. The SNARE complex is formed by the interaction of plasma membrane-anchored proteins Syntaxin-1 (STX1 refers to Syntaxin-1A and Syntaxin-1B throughout this study unless stated otherwise) and SNAP-25 and vesicle-associated membrane protein Synaptobrevin-2 (VAMP/SYB2). The SNARE proteins assemble their 55 amino acid-long, highly conserved SNARE motifs into a parallel 4-helix bundle. The resulting SNARE complex that zippers up sequentially from N-to C-terminus physically brings the vesicle and plasma membranes together and provides the necessary energy to mechanically force membrane merger (Fasshauer et al., 1998; Sutton et al., 1998; Sørensen et al., 2006; Weber et al., 2010; Zhang et al., 2017, 2022). The assembly of the SNARE complex creates a highly charged outer surface (Fasshauer et al., 1998; Sutton et al., 1998; Ruiter et al., 2019) to which accessory proteins such as complexins, synaptotagmins or Sec1/Munc18(SM)-proteins can bind to and thereby modulate the efficacy and timing of neurotransmitter release (Rizo, 2022; Rizo & Rosenmund, 2008; Südhof, 2013).

Among the unique components of the SNARE complex, STX1 stands out by its complex structural characteristics. On the N-terminal extreme of STX1, the short regulatory N-peptide and the autonomously folded three-helical H_abc_-domain are essential for the formation of fusogenic SNARE complexes and the readily releasable pool (RRP) of vesicles (Dulubova et al., 2007; Fernandez et al., 1998; Gerber et al., 2008; Khvotchev et al., 2007; Ma et al., 2011; Vardar et al., 2021; Zhou et al., 2013). On the C-terminal extreme, STX1 contains the juxtamembrane domain (JMD) and the transmembrane domain (TMD) that are actively involved in membrane fusion (Dhara et al., 2016; Gao et al., 2012; McNew et al., 1999; Sutton et al., 1998; Vardar et al., 2022). The most crucial, however, is its SNARE domain as it directly regulates the membrane merger. SNAP-25 and SYB2 mutation studies that destabilize the inner hydrophobic pocket of the SNARE domain have determined that the SNARE motif of these proteins can be functionally compartmentalized into two subdomains: the N-terminus, necessary for priming and nucleation of the SNARE complex, and the C-terminus, essential for fusion (Gao et al., 2012; Sørensen et al., 2006; Walter et al., 2010; Weber et al., 2010). This was further confirmed by single-molecule optical tweezer studies which determined the sequential assembly of the SNARE complex through diverse staged assisted by these two subdomains (Gao et al., 2012). Additionally, it has been determined that accessory proteins, such as Munc18-1 and Complexin, chaperone the stability of the N-terminus and facilitate or clamp the assembly of the C-terminus in the partially-zippered SNARE domain, which without their regulation is fast and spontaneous (Gao et al., 2012; Hao et al., 2023; Ma et al., 2015). Although studies using STX1/STX3 chimeric constructs revealed that the SNARE motif of STX1 may carry structural features that regulate spontaneous release (Vardar et al., 2022), research on the SNARE domain of STX1 has been limited to the studies of its cognate interaction partners such as Munc18-1, Synaptotagmin-1 (SYT1) and Complexin (Hao et al., 2023; Jiao et al., 2018; Ma et al., 2015; Zhou et al., 2015, 2017).

In this study, we address the specific function of the entire SNARE domain of STX1 in regulating neurotransmitter release in intact synapses. For that purpose, we used a comprehensive structure-function approach and compared the efficacy of STX1 to the closely-related isoform STX2 (Sherry et al., 2006), which is implicated in the secretion of various peripheral secretory cells (Abonyo et al., 2004; Dolai et al., 2018; Hutt et al., 2005). When expressed in STX1-null hippocampal mouse neurons (Vardar et al., 2016), STX2 synapses showed impaired vesicle priming, unclamping of spontaneous release, as well as inefficient and slowed Ca^2+^-evoked fusion, consistent with an impaired SNARE complex function (Hao et al., 2023; Jiao et al., 2018; Lai et al., 2017; Stepien et al., 2022; Voleti et al., 2020). Therefore, we proceeded with a structure-function chimeric analysis between STX1 and STX2, to isolate the precise involvement of the SNARE domain of STX1 in the regulatory mechanisms of release. We found that the C-terminal half of the SNARE domain of STX1 is a key regulator in the stability of the readily releasable pool and is part of the clamping mechanism for spontaneous release in central synapses. Additionally, while it may have a role in the regulation of the speed and efficacy of Ca^2+^-evoked release, these functions depend on the integrity of the full SNARE domain and other domains of STX1. Overall, we discovered that the role of the SNARE complex goes beyond zippering and interacting with regulators of release and has evolved to participate in the finetuning of the fusion energy barrier and the precision of synaptic vesicle release alongside its accessory proteins.

## Results

### STX2 supports neuronal viability

Previously we have shown that comparison of non-canonical syntaxin isoforms to STX1 can offer insight into the specificity of vesicle fusion (Vardar et al., 2022). For this, Syntaxin-2 (STX2), Syntaxin-3 (STX3) and Syntaxin-4 (STX4) are the best candidates as they share the same domain structure with STX1 (Quiñones et al., 1999). For example, the isoform STX2 has been found in brain tissue and is present in the synaptosome fraction (Chen et al., 1999). STX1A and STX2 share a 63% total sequence homology and a 69% homology in their SNARE domain (Figure 1A). As expected from the promiscuity of syntaxins in particular and SNARE proteins in general, overexpression of STX2 can rescue neuronal survival in the absence of functional STX1 by binding to SNAP-25 (Peng et al., 2013), although its cognate SNARE partner in non-neuronal intracellular trafficking is SNAP-23 (Abonyo et al., 2004).

**Figure 1.**
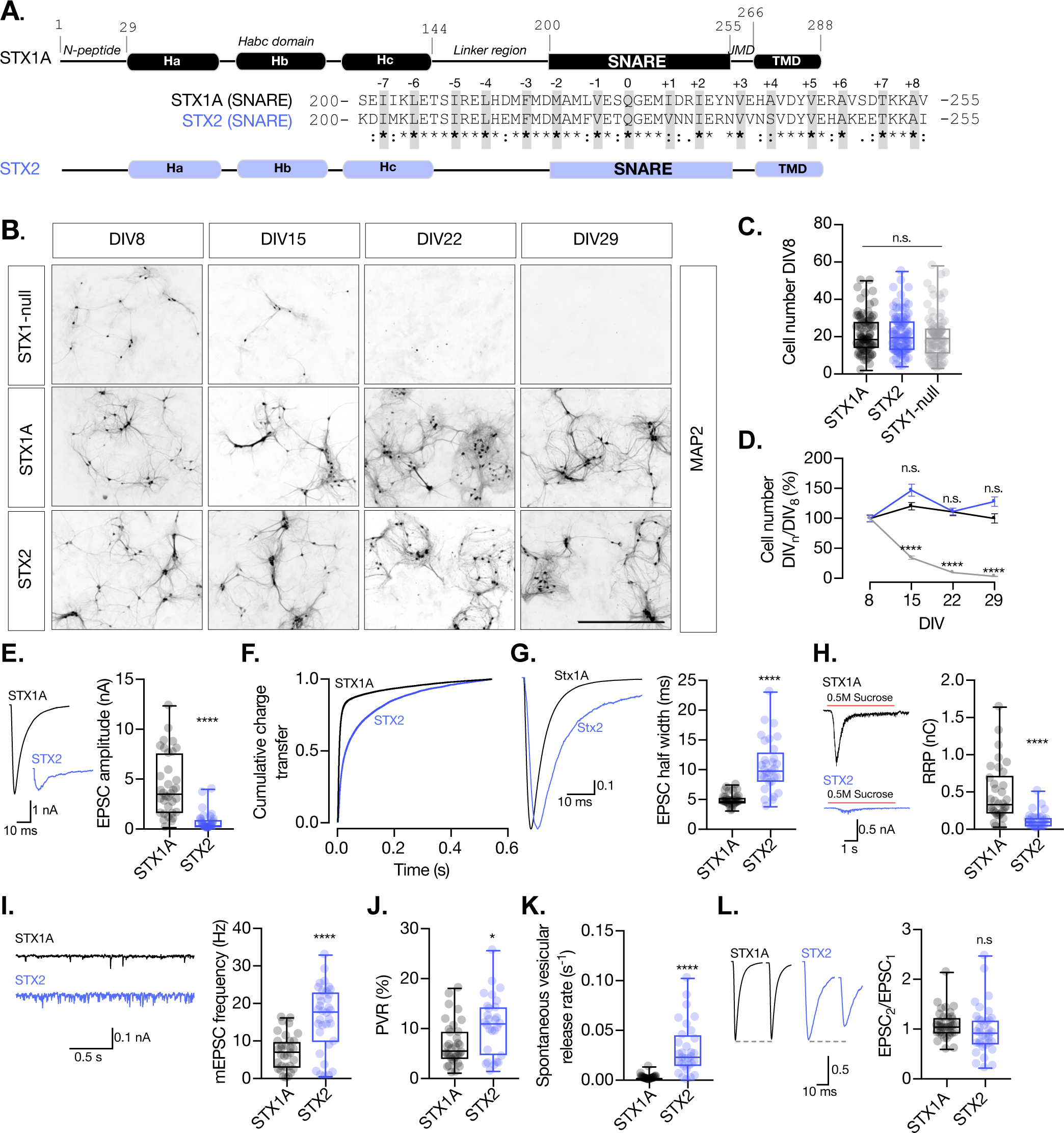
STX2 supports neuronal viability but does not rescue synchronous evoked release, the RRP or the clamping of spontaneous release. A. STX1A and STX2 domain structure scheme and SNARE domain sequence alignment (68% homology). Layers are highlighted in gray. B. Example images of high-densitiy cultured STX1-null hippocampal neurons rescued with GFP (STX1-null), STX1A or STX2 (from top to bottom) at DIV8, DIV15, DIV22 and DIV29 (from left to right). Immunofluorescent labelling of MAP2. Scale bar: 500μm. C. Quantification of total number of neurons at DIV8 of each group. D. Quantification of the percentage of the surviving neurons at DIV8, DIV15, DIV22 and DIV29 normalized to the number of neurons at DIV8 in the same group. E. Example traces (left) and quantification of EPSC amplitude (right) from autaptic Stx1A-null hippocampal mouse neurons rescued with STX1A or STX2. F. Quantification of the cumulative charge transfer of the EPSC from the onset of the response until 0.55s after. G. Example traces of normalized EPSC to their peak amplitude (left) and quantification of the half width of the EPSC (right). H. Example traces (left) and quantification of the response induced by a 5s 0.5M application of sucrose, which represents the readily releasable pool of vesicles (RRP). I. Example traces (left) and quantification of the frequency of the miniature excitatory postsynaptic currents (mEPSC) (rights). J. Quantification of the vesicle release probability (PVR) as the ratio of the EPSC charge over the RRP charge (PVR). K. Quantification of the spontaneous vesicular release rate as the ratio between the of mEPSC frequency and number of vesicles in the RRP. L. Example traces (left) and the quantification of a 40Hz paired-pulse ratio (PPR). Data information: In (D) data points represent mean ± SEM. In (D, E-L) data is show in a whisker-box plot. Each data point represents single observations, middle line represents the median, boxes represent the distribution of the data and external data points represent outliers. In (C, D) significances and P values of data were determined by non-parametric Kruskal-Wallis test followed by Dunn’s post hoc test; *p≤0.05, **p≤0.001, ***p≤0.001, ****p≤0.0001. In (E-L) Significances and P values of data were determined by non-parametric Mann-Whitney test and unpaired two-tailed t-test; *p≤0.05, **p≤0.01, ***p≤0.001, ****p≤0.0001. All data values are summarized in Figure 1 – Source Data 1.

Having this in mind, we lentivirally expressed STX2 in STX1-null neurons using our well-established model of STX1-null mouse hippocampal neurons (Vardar et al., 2016) and conducted a series of rescue experiments. STX1-null neurons typically do not survive in culture longer that 8 days *in vitro* (DIV) and exhibit a complete loss of neurotransmitter release (Vardar et al., 2016) (Figure 1C). Yet healthy synapses are vital to provide a solid background to further analyze the electrophysiological properties. Therefore, our first objective was to compare the ability of STX1A and STX2 isoforms to rescue neuronal survival. We quantified the amount of mouse hippocampal neurons in high-density neuronal cultures at DIV8, DIV15, DIV22 and DIV29 for each group and found that STX2 rescues neuronal survival at a comparable level to STX1A (Fig. 1B and 1D). Furthermore, immunocytochemistry experiments confirmed that STX2, as well as STX1A, is strongly expressed in our model neurons (Figure 1-figure supplement 1).

### STX2 does not support synchronous Ca^2+^-evoked release, the RRP size, nor the spontaneous release <u>clamp.</u>

Next, we analyzed the electrophysiological properties of autaptic neurons rescued with STX1A or STX2. First, STX2 neurons showed a dramatic reduction in the excitatory postsynaptic current (EPSC) amplitude from 4.3nA (SEM±0.53) to 0.7nA (SEM±0.16) (Figure 1E). We plotted the cumulative charge transfer of the EPSC over more than 550ms after the stimulation. A rapid and synchronous release of vesicles was reflected in a much more rapid cumulative charge transfer in STX1A neurons (black line; Figure 1F), compared to STX2 neurons, where the cumulative charge transfer of the response was much slower (purple line; Figure 1F). Additionally, the half width of the EPSC was increased more than 2-fold in STX2 neurons (Figure 1G). This resonates with previous studies that showed a decrease in the speed of secretion of STX2-expressing non-neuronal cells (Abonyo et al., 2004). Overall, our results indicate that STX2 does not support synchronous Ca^2+^-evoked release, but a slow asynchronous response. We also evaluated synaptic vesicle priming by applying a hypertonic sucrose solution (500mM) for 5s and measuring the charge of the response (Rosenmund & Stevens, 1996) and found that STX2 neurons exhibit a 75% reduction in the readily releasable pool (RRP) of vesicles compared to STX1A neurons (Figure 1H). This suggests that STX2 does not support synaptic vesicle priming to the same extent as STX1A. STX2-expressing neurons also displayed more than 2-fold increase in the frequency of spontaneous release (mEPSC) relative to STX1A neurons (Figure 1I) and a 55% increase in the probability of release (PVR), calculated by dividing the EPSC charge by the RRP charge (Figure 1J). When we normalized the mEPSC frequency to the RRP size we found a nearly 15-fold increase in the spontaneous vesicular release rate in STX2 compared to STX1A neurons (Figure 1K). Finally, no change was observed in the paired-pulse ratio (PPR) between STX1A and STX2 groups (Figure 1L). Notably, the difficulty of analysis given the small values of the release of the second pulse for STX2 neurons makes it hard to interpret this data. The increase in the spontaneous vesicular release rate and the PVR indicates that STX2-containing SNARE complexes render vesicles more fusogenic. Taken together, this data suggests that STX2 neurons follow a fusion energetic landscape that differs from STX1A and resembles constitutive versus regulated release (reviewed by Sørensen, 2009).

### The C-terminal half of the SNARE domain of STX1A has a regulatory effect on the RRP size and on spontaneous release

Previous structure-function studies have shown the critical role of the distinct domains of STX1 in different aspects of synaptic vesicle fusion. However, what characterizes the protein is its SNARE domain. Not only does the SNARE domain of STX1A form one of the 4 helixes of the SNARE complex (Sutton et al., 1998), it also interacts with several regulatory proteins that modulate priming and fusion, such as SM protein Munc18-1, calcium sensor Synaototagmin-1 (SYT1) and accessory protein Complexin. Additionally, the SNARE domain has been compartmentalized into two functional subdomains: N-terminus, key in priming the vesicles to the plasma membrane and C-terminus, critical for the fusion process (Gao et al., 2012; Schotten et al., 2015; Sørensen et al., 2006; Walter et al, 2010; Weber et al., 2010). Our neuronal survival test shows that we can utilize STX2 for electrophysiological studies. To confine our studies to the SNARE domain, we used a chimeric approach, where we exchanged either the entire SNARE domain or the N-or C-terminal halves of the SNARE domain (N-terminal from E/D200-Q226; C-terminal from Q226-V/I255) between STX1A and STX2 (Figure 2A). Whereas the N-terminal halves of STX1A and STX2 share an 81% homology, their C-terminal halves, only share 60% homology (Figure 2A), and each half potentially contributes to a different step in vesicle fusion (Gao et al., 2012; Sørensen et al., 2006; Weber et al., 2010).

**Figure 2.**
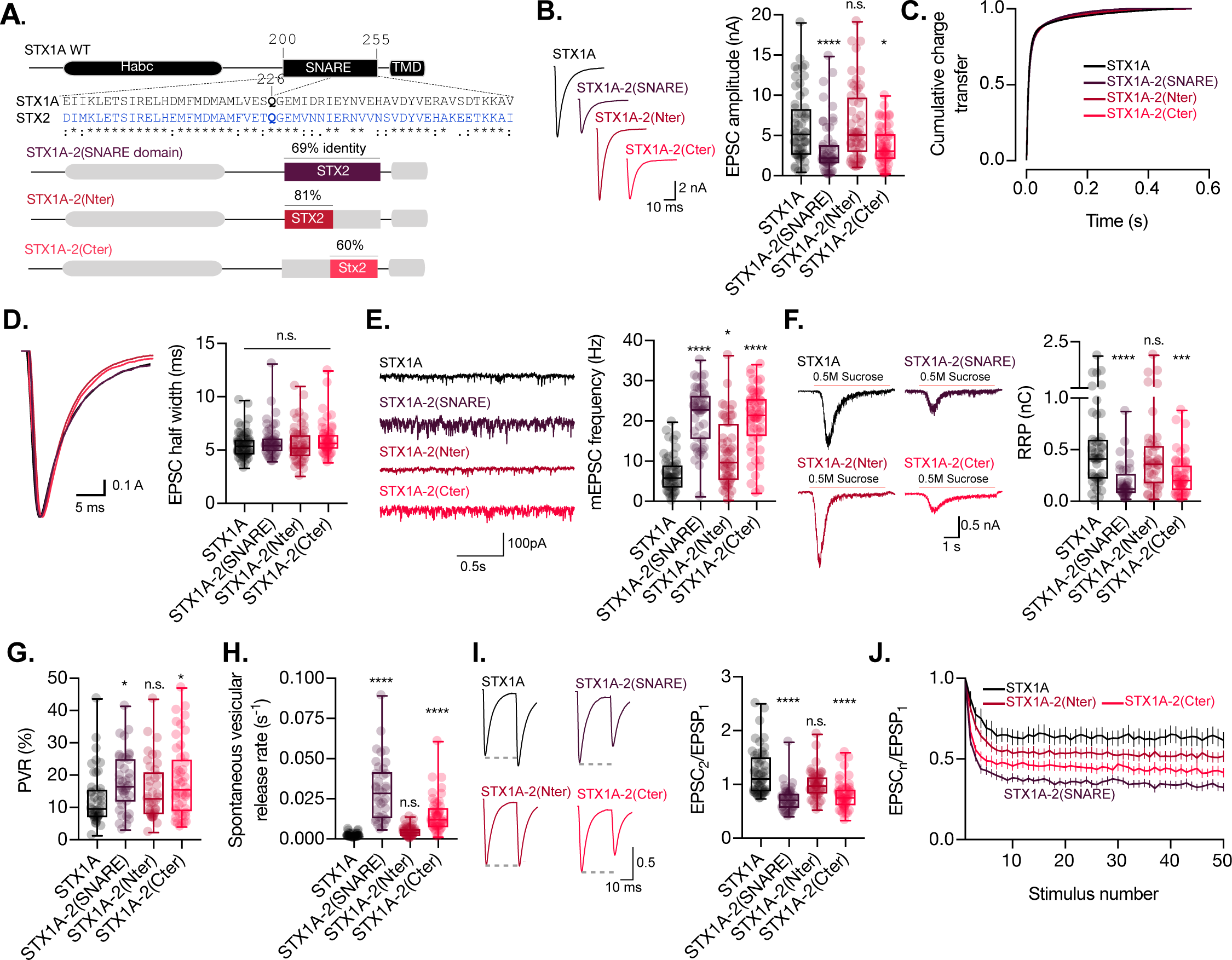
The C-terminal half of the SNARE domain of STX1A has a regulatory effect on the RRP, spontaneous release and both, N- and C-terminus have a role in the regulation of efficacy of Ca^2+^-evoked release. A. STX1A WT and chimera domain structure scheme, sequence alignment of STX1A and STX2 SNARE domain and percentage homology between both SNARE domains. B. Example traces (left) and quantification of the EPSC amplitude (right) from autaptic STX1A-null hippocampal mouse neurons rescued with STX1A, STX1A-2(SNARE), STX1A-2(Nter) or STX1A-2(Cter). C. Quantification of the cumulative charge transfer of the EPSC from the onset of the response until 0.55s after. D. Example traces of normalized EPSC to their peak amplitude (left) and quantification of the half width of the EPSC (right). E. Example traces (left) and quantification of the frequency of the miniature excitatory postsynaptic currents (mEPSC) (right). F. Example traces (left) and quantification of the response induced by a 5s 0.5M application of sucrose, which represents the readily releasable pool of vesicles (RRP). G. Quantification of the vesicle release probability (PVR) as the ratio of the EPSC charge over the RRP charge (PVR). H. Quantification of the spontaneous vesicular release rate as the ratio between the of mEPSC frequency and number of vesicles in the RRP. I. Example traces (left) and the quantification of a 40Hz paired-pulse ratio (PPR). J. Quantification of STP measured by 50 stimulations at 10Hz. Data information: In (B, D-I) data is show in a whisker-box plot. Each data point represents single observations, middle line represents the median, boxes represent the distribution of the data, where the majority of the data points lie and external data points represent outliers. In (J) data represents the mean ± SEM. Significances and P values of data were determined by non-parametric Kruskal-Wallis test followed by Dunn’s post hoc test; *p≤0.05, **p≤0.01, ***p≤0.001, ****p≤0.0001. All data values are summarized in Figure 1 – Source Data 2.

Replacement of the full-SNARE domain (STX1A-2(SNARE)) or the C-terminal half (STX1A-2(Cter)) of the SNARE domain of STX1A with the same domain from STX2 resulted in a reduction in the EPSC amplitude (Figure 2B). However, there was no change in the cumulative charge transfer (Figure 2C) or the kinetic parameters of the EPCS, such as half width (Figure 2D) or rise and decay time (Figure 2-figure supplement 1A and B), in any of the groups. This suggests that the SNARE domain of STX1A may not be directly involved in the regulation of the kinetics of the release. Our analysis also showed that neurons expressing STX1A-2(SNARE) or STX1A-2(Cter) exhibited a nearly 3-fold increase in the mEPSC frequency (Figure 2E), potentially indicating an unclamping of spontaneous release. Additionally, the total charge transfer in response to a hypertonic sucrose application was significantly decreased in these two groups from 0.5nC (SEM±0.06) to 0.18nC (SEM±0.02) and 0.2nC (SEM±0.02) respectively, compared to STX1A WT, indicating fewer primed synaptic vesicles and a smaller RRP (Figure 2F). Notably, the common modification of these STX1A chimera groups is the addition of the C-terminal half of the SNARE domain of STX2. These results suggest that the C-terminal half of the SNARE domain of STX1A may be directly involved in the clamping mechanism for spontaneous release of synaptic vesicles and may also play a role in the modulation of the RRP. However, it should be noted that we also observed a significant increase from 6.84Hz (SEM±0.62) to 11.8Hz (SEM±1.28) in the mEPSC frequency of STX1A-2(Nter)-expressing neurons (Figure 2E). Therefore, we cannot rule out the possibility that the N-terminal half of the SNARE domain may also play a role in the regulation of spontaneous release.

We next examined release efficacy in the STX1A-STX2 SNARE domain chimeras. STX1A-2(SNARE) showed a 39% increase and STX1A-2(Cter) showed a 42% increase in the PVR (Figure 2G), which indicates that the reduction in the EPSC is not proportional to the decrease in the RRP. This suggests that the changes we observed could be due to a faulty priming mechanism, but it could also mean that for these two groups there is a change in an additional mechanism that goes beyond the RRP size regulation and increases the efficacy of the Ca^2+^-evoked release. Finally, STX1A-2(SNARE) and STX1A-2(Cter) had a 17- and 8-fold increase, respectively, in the spontaneous vesicular release rate (Figure 2H), a 40% decrease in the pair-pulse ratio (PPR) (Figure 2I) and increased depression in a 10Hz train stimulation (Figure 2J). Taken together our results suggest that the C-terminal half of the SNARE domain of STX1A is involved in the regulation of the efficacy of Ca^2+^-evoked release, the formation of the RRP and in the clamping of spontaneous release.

### The insertion of the full-length SNARE domain or the C-terminal half of the SNARE domain of STX1A into the STX2 backbone has an impact on the kinetics of evoked release, RRP size and spontaneous release

In the same manner as before, we wanted to test whether any of the electrophysiological properties that differ between the two syntaxin isoforms are attributable to the SNARE domain itself and can be transferred to STX2. For this, we generated chimeric proteins by introducing the entire SNARE domain of STX1A (STX2-1A(SNARE)), its N-terminal half (STX2-1A(Nter)) or its C-terminal half (STX2-1A(Cter)) into STX2 (Figure 3A). Not only does this chimeric approach allow us to isolate certain functions of the SNARE domain, but it can also help us determine whether the function of the SNARE domain of STX1A depends on domains or sequences present in its natural structure.

**Figure 3.**
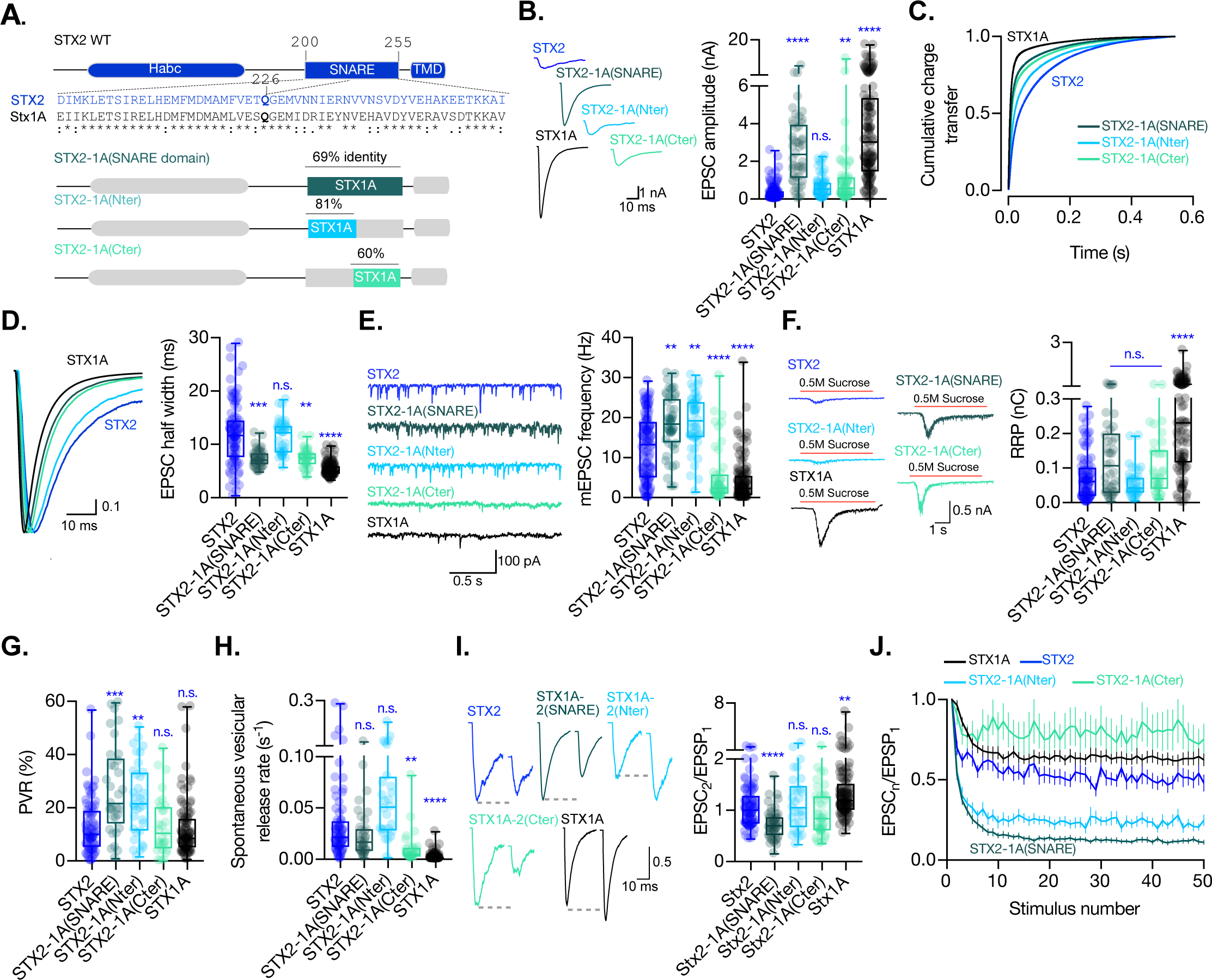
The C-terminal half of the SNARE domain of STX1A has a regulatory effect on the spontaneous release and the RRP and the speed of Ca^2+^-evoked release depends on the integrity of the SNARE domain. A. STX2 WT and chimera domain structure scheme, sequence alignment of STX2 and STX1A SNARE domain and percentage homology between both SNARE domains. B. Example traces (left) and quantification of the EPSC amplitude (right) from autaptic STX1A-null hippocampal mouse neurons rescued with STX2, STX2-1A(SNARE), STX2-1A(Nter) or STX2-1A(Cter). C. Quantification of the cumulative charge transfer of the EPSC from the onset of the response until 0.55s after. D. Example traces of normalized EPSC to their peak amplitude (left) and quantification of the half width of the EPSC (right). E. Example traces (left) and quantification of the frequency of the miniature excitatory postsynaptic currents (mEPSC) (right). F. Example traces (left) and quantification of the response induced by a 5s 0.5mM application of sucrose, which represents the readily releasable pool of vesicles (RRP). G. Quantification of the vesicle release probability (PVR) as the ratio of the EPSC charge over the RRP charge (PVR). H. Quantification of the spontaneous vesicular release rate as the ratio between the of mEPSC frequency and number of vesicles in the RRP. I. Example traces (left) and the quantification of a 40Hz paired-pulse ratio (PPR). J. Quantification of STP measured by 50 stimulations at 10Hz. Data Information: In (E-L) data is show in a whisker-box plot. Each data point represents single observations, middle line represents the median, boxes represent the distribution of the data, where the majority of the data points lie and external data points represent outliers. Significances and P values of data were determined by non-parametric Kruskal-Wallis test followed by Dunn’s post hoc test; *p≤0.05, **p≤0.01, ***p≤0.001, ****p≤0.0001. All data values are summarized in Figure 3 – Source Data 1.

When recording from the STX2-STX1A SNARE domain chimera-expressing neurons we observed that we were able to partially rescue the EPSC amplitude when STX2 contained the entire SNARE domain of STX1A (STX2-1A(SNARE)) compared to STX2, from 0.35nA (SEM±0.044) to 2.6nA (SEM±0.28), respectively, although not to STX1A WT levels 4.06nA (SEM±0.31). Additionally, STX2-1A(Cter) showed a slight, albeit significant, rescue of the amplitude of the Ca^2+^-evoked response to 1.13nA (SEM±0.28). STX2-1A(Nter) also showed a 50% increase in the Ca^2+^-evoked release to 0.68nA (SEM±0.08) (Figure 3B), suggesting an insufficient rescue of the evoked response. To analyze the kinetics of release in STX2-chimeras, we plotted the cumulative charge transfer of the EPSC over time for each group and quantified kinetic parameters such as half width, rise time and decay time. STX2 and STX2-1A(Nter) exhibited the slowest cumulative charge transfer (Figure 3C, blue and light blue line, respectively) as well as the highest half width (Figure 3D) and rise and decay time (Figure 3-figure supplement 1A and B). These findings indicate that the N-terminal half of the SNARE domain of STX1A is not enough to support the fast kinetics of the response. However, the introduction of the entire SNARE domain or the C-terminal half rescued the speed and synchronicity of the EPSC to almost STX1A WT levels (Figure 3C, dark and light turquois lines, respectively, Figure 3D and Figure 3-figure supplement 1A and B). Notably, the speed of release did not change in any STX1A-chimera (Figure 2C and D), suggesting that there is a dominant regulatory domain which supports the fast kinetics of the response outside of the SNARE domain in STX1A, such as the juxtamembrane or transmembrane domain (Vardar et al., 2022). However, the C-terminal half of STX1A alone is sufficient to evoke faster responses in STX2. These results support major regulatory differences of the domains outside of the SNARE domain between the isoforms, which should not be discarded in the interpretation of our results.

As reported previously, our STX1A chimeric analysis suggested a clamping function of the C-terminal half of the SNARE domain of STX1A (Figure 2E). In line with these findings, STX2-1A(Cter) showed a significant reduction in the mEPSC frequency to almost STX1A WT levels. STX2-1A(SNARE) unexpectedly showed a 47% increase in the spontaneous release frequency compared to STX2, which was also observed in STX2-1A(Nter) (Figure 3E). Additionally, we found that STX2-1A(SNARE) and STX2-1A(Cter) could rescue the RRP to around double of what we measured from STX2 and STX2-1A(Nter) (Figure 3F), albeit not significant. This trend supports our hypothesis that the C-terminal half of the SNARE domain of STX1A potentially plays a role in the regulation in vesicle priming, also revealed by STX1A-chimeras (Figure 2F). Furthermore, STX2-1A(SNARE) and STX2-1A(Nter) had an increased PVR (Figure 3G), no change in the release rate (Figure 3H) and an increase in short-term depression during 10Hz train stimulation (Figure 3J), while only STX2-1A(SNARE) in the PPR (Figure 3I), compared to STX2 WT. On the other hand, STX2-1A(Cter) showed no change in the PVR compared to STX2 or STX1A WT, a 4.5-fold decrease in the spontaneous release rate (STX1A WT has a 12-fold decrease compared to STX2), and an increase in the short-term depression in a 10Hz train stimulation albeit the difficulty of analysis of this data given the small values of the release (Figure 3J). These data indicate that the suppression of spontaneous release and the regulation of the RRP relies on the C-terminal half of the SNARE domain of STX1A, while the efficacy of Ca^2+^-evoked release depends on the integrity of the entire SNARE domain (Figure 3G, 3I and 3J).

### The insertion of the full-length SNARE domain or the C-terminal half of the SNARE domain of STX1A into the Stx2 backbone rescues Munc18-1 levels at the synapse

Munc18-1 binds not only to the N-terminal regions of STX1A (Burkhardt, Hattendorf, Weis, & Fasshauer, 2008; Khvotchev et al., 2007; Meijer et al., 2018) but also to the SNARE complex (Baker et al., 2015; Jiao et al., 2018; Stepien et al., 2022). The expression levels of Munc18-1 and STX1A influence each other and their interaction is important for priming and release (Gerber et al., 2008; Vardar et al., 2016). Thus, we analyzed the exogenous expression of STX1A, STX1A-2(SNARE), STX1A-2(Nter) and STX1A-2(Cter) and the endogenous expression of Munc18-1 at the synapse (Figure 4). We used VGlut1 as the synaptic marker in the co-localization assays. We found that the introduction of the SNARE domain of STX2 into STX1A (STX1A-2(SNARE)) increased the expression of this construct compared to STX1A WT while the rest showed similar levels (Figure 4B). Additionally, the exchange of the entire SNARE domain or the C-terminus of STX1A with that of STX2 caused an increase in the levels of endogenous Munc18-1 at the synapse, which may be relative to the increase in STX1A-construct levels (Figure 4C). Because the loss or overexpression of Munc18-1 affect the properties of secretion and release (Oh et al., 2012; Toonen et al., 2006; Verhage et al., 2000; Voets et al., 2001), we wanted to determine whether changes in the levels of Munc18-1 may correlate with the changes in neurotransmitter release from the neurons which carry the exogenous STX1A-chimeric constructs. We observed that higher levels of Munc18-1 seem to correlate with less Ca^2+^-evoked release (Figure 4E), and a smaller RRP (Figure 4F) but higher release efficacy (Figure 4G) and release rates (Figure 4H), which suggest it may be the cause of these changes in the release parameters.

**Figure 4.**
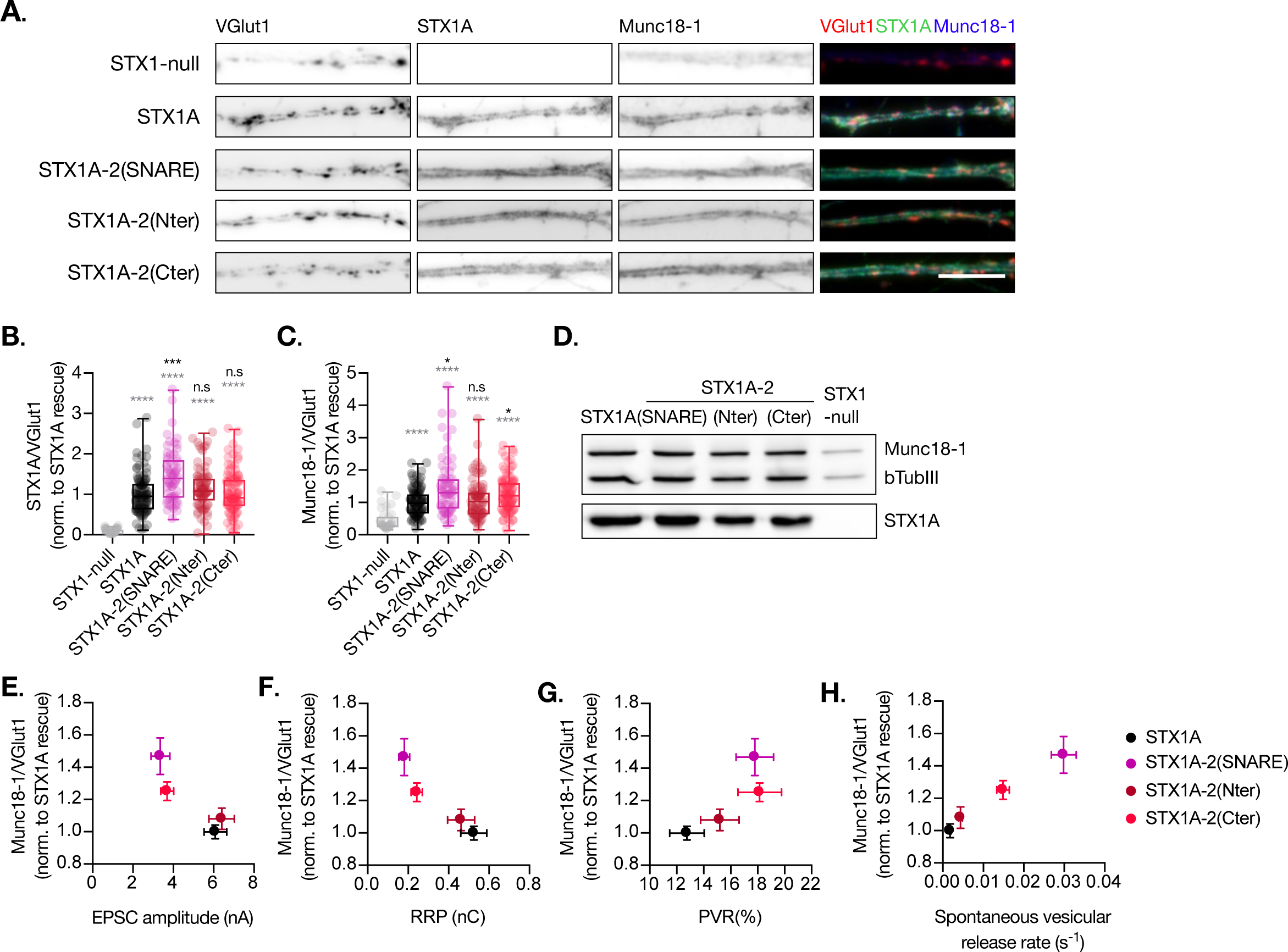
Quantification of STX1A and Munc18-1 levels at the synapse. A. Example images of Stx1-null neurons plated in high-density cultures and rescued with STX1A, STX1A-2(SNARE), STX1A-2(Nter) or STX1A-2(Cter) or GFP (STX1-null) as negative control. Neurons were fixed between DIV14-16. Cultures were stained with fluorophore-labeled antibodies that recognize VGlut1 (red in merge), STX1A (green in merge) and Munc18-1(blue in merge), from left to right. Scale bar: 10μm. B. Quantification of the immunofluorescent intensity of STX1A normalized to the intensity of the same VGlut1-labeled ROIs. Values where normalized to values in STX1A rescue group. C. Quantification of the immunofluorescent intensity of Munc18-1 normalized to the intensity of the same VGlut1-labeled ROIs. Values where normalized to values in STX1 rescue group. D. SDS-PAGE of the electrophoretic analysis of neuronal lysates obtained from each experimental group. Proteins were detected using antibodies that recognize β-Tubuline III as loading control, STX1A and Munc18-1. E. Correlation between Munc18-1 values and EPSC amplitude of Stx1-null neurons expressing STX1A, STX2, STX2-1A(SNARE), STX2-1A(Nter) or STX2-1A(Cter). F. Correlation between Munc18-1 values and RRP. G. Correlation between Munc18-1 values and PVR. H. Correlation between Munc18-1 values and spontaneous vesicular release rate Data information: In (B, C) data is show in a whisker-box plot. Each data point represents single ROIs, middle line represents the median, boxes represent the distribution of the data, where the majority of the data points lie and external data points represent outliers. In (E-H) each data point is the correlation of the mean ±SEM. Significances and P values of data were determined by non-parametric Kruskal-Wallis test followed by Dunn’s post hoc test; *p≤0.05, **p≤0.01, ***p≤0.001, ****p≤0.0001. All data values are summarized in Figure 4 – Source Data 1.

Additionally, we quantified the levels of exogenous expression of STX2, STX2-1A(SNARE), STX2-1A(Nter) and STX2-1A(Cter) and the corresponding levels of endogenous Munc18-1 levels at the synapse (Figure 5). We found that all STX2 constructs were expressed at similar levels at the synapse (co-localized to VGlut1 puncta) (Figure 5A and B). However, STX2-1A(SNARE) and STX2-1A(Cter) showed a trend towards increase in their expression (Figure 5B and C). We then quantified the endogenous levels of Munc18-1 at the synapse in these neurons and found a decrease in Munc18-1 levels when expressing STX2 WT compared to STX1A WT. Interestingly, Munc18-1 levels were rescued when either the entire SNARE domain (STX2-1A(SNARE) or the C-terminal half (STX2-1A(Cter)) of STX1A were present (Figure 5E and 5F) compared to STX1A WT neurons. The interaction between Munc18-1 and the SNARE domain, which is thought to be important for templating SNARE complex assembly (Baker et al., 2015; Stepien et al., 2022), may be directly involved in rescuing some of the electrophysiological parameters observed in STX2 chimeras. To investigate this, we correlated the levels of Munc18-1 with electrophysiological phenotypes observed in neurons expressing the STX2-chimeras (Figure 5G-J). Our superposition of phenotypes revealed that higher levels of Munc18-1, observed in STX1A WT and in the STX2-1A(Cter) neurons, showed a trend in increasing Ca^2+^-evoked release (Figure 5G) and the RRP (Figure 5H) while the effects on PVR were more randomly distributed (Figure 5I). Additionally, we observed that higher release rates correspond to groups which show less amounts of Munc18-1 at the synapse (Figure 5J). Taking together these results it seems like we have contradictory observations: while increased levels of Munc18-1 in the STX1A background seem to correlate with parameters that depict an increase in the fusogenicity of the vesicles (Figure 4), an increase in the levels of Munc18-1 in the STX2 background correlates to release properties that incline towards a decreased fusogenicity. In summary, although Munc18-1 levels does depend on the construct used and may be causative of changes in certain electrophysiological characteristics, we may not be able to explain these phenotypes by the levels of Munc18-1.

**Figure 5.**
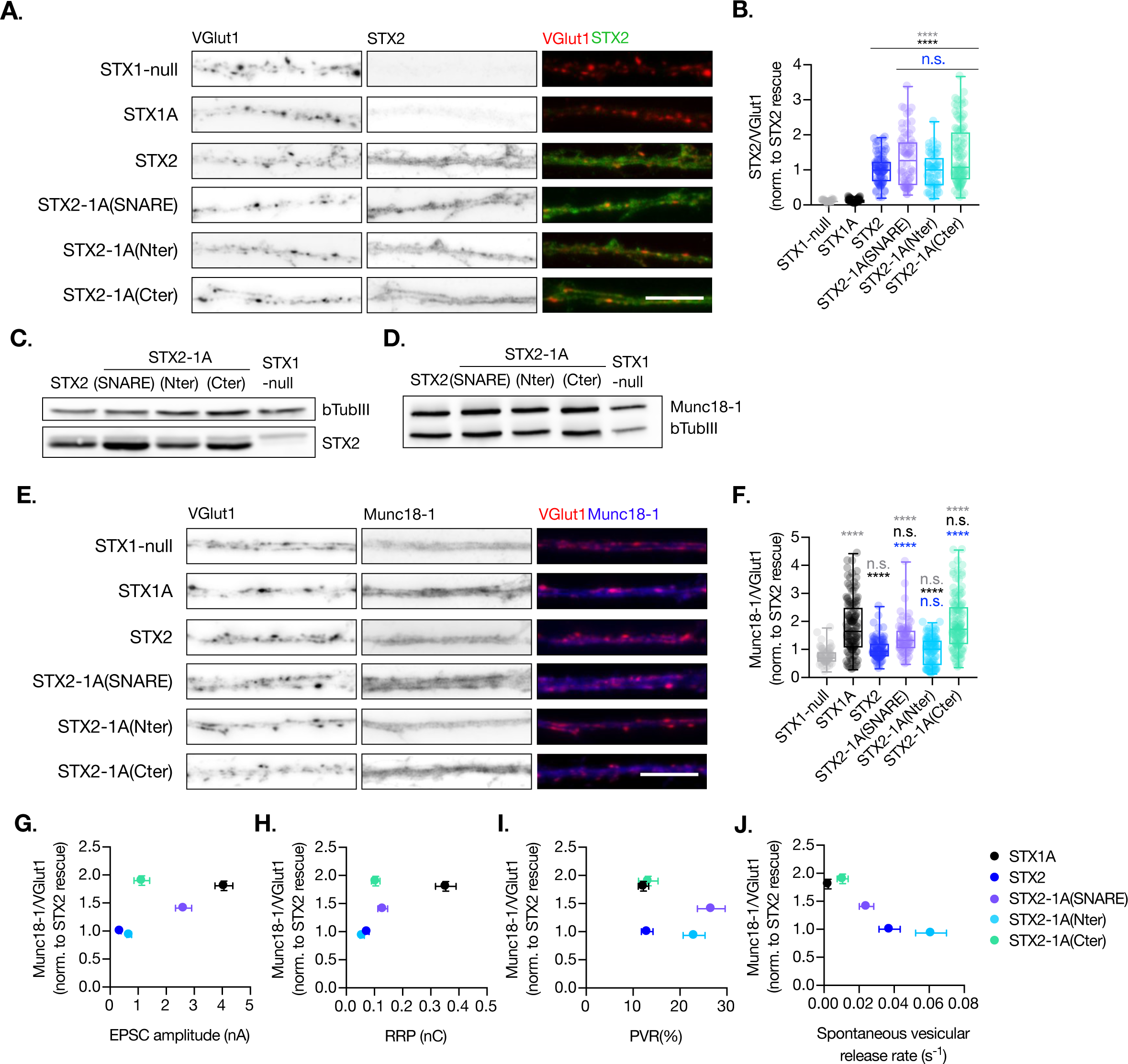
Quantification of STX2 and Munc18-1 levels at the synapse. A. Example images of Stx1-null neurons plated in high-density cultures and rescued with STX1A, STX2, STX2-1A(SNARE), STX2-1A(Nter), STX2-1A(Cter) or GFP (STX1-null) as negative control. Between DIV14-16 neurons were fixed**. (A)** Example images of neurons stained with fluorophore-labeled antibodies that recognize VGlut1 (red in merge) and STX2 (green in merge), from left to right. Scale bar: 10μm. B. Quantification of the immunofluorescent intensity of STX2 normalized to the intensity of the same VGlut1-labeled ROIs. Values where normalized to values in STX2 rescue group. C. SDS-PAGE of the electrophoretic analysis of neuronal lysates obtained from each experimental group. Proteins were detected using antibodies that recognize β-Tubuline III as loading control and STX2. D. SDS-PAGE of the electrophoretic analysis of neuronal lysates obtained from each experimental group Proteins were detected using antibodies that recognize β-Tubuline III and Munc18-1. E. Example images of neurons stained with fluorophore-labeled antibodies that recognize VGlut1 (red in merge) and Munc18-1(blue in merge), from left to right. Scale bar: 10μm. F. Quantification of the immunofluorescent intensity of Munc18-1 normalized to the intensity of the same VGlut1-labeled ROIs. Values where normalized to values in STX2 rescue group. G. Correlation between Munc18-1 values and EPSC amplitude of STX1-null neurons expressing STX1A, STX2, STX2-1A(SNARE), STX2-1A(Nter) or STX2-1A(Cter). H. Correlation between Munc18-1 values and RRP. I. Correlation between Munc18-1 values and PVR. J. Correlation between Munc18-1 values and spontaneous vesicular release rate. Data information: In (B, F) data is show in a whisker-box plot. Each data point represents single ROIs, middle line represents the median, boxes represent the distribution of the data, where the majority of the data points lie and external data points represent outliers. In (G-J) each data point is the correlation of the mean ±SEM. Significances and P values of data were determined by non-parametric Kruskal-Wallis test followed by Dunn’s post hoc test; *p≤0.05, **p≤0.01, ***p≤0.001, ****p≤0.0001. All data values are summarized in Figure 5 – Source Data 1.

### The residues on the outer surface of the C-terminal half of the SNARE domain are crucial in the regulation of spontaneous release and the RRP

So far, we have highlighted the importance of the C-terminal half of the SNARE domain of STX1A in the regulation of the RRP size and for the clamping of spontaneous release. Our previous study that aligned STX1A and different STX3A and STX3B chimeric constructs also suggested the involvement of the C-terminus half in the spontaneous release clamping mechanisms, but only through elimination (Vardar et al., 2022). These data compelled us to examine whether spontaneous release regulatory mechanisms can be attributed to individual or sequentially paired amino acid residues in the C-terminal half of the SNARE domain of STX1A. To test this, we introduced single-or double-point mutations in the C-terminus of the SNARE domain of STX1A, targeting residues that differ from STX2 in their charge or polarity and that are located on the outer surface of the SNARE complex. It is important to note that electrostatic charges play a critical role in the interaction of STX1A‘s SNARE domain with other proteins and the membrane (Ruiter et al., 2019). Thus, we hypothesized that if the charge of the amino acid was not significantly altered, it may not play a significant role in the observed phenotypic differences between STX1A and STX2. Given this we generated single point mutations (STX1A^D231N^, STX1A^R232N^, STX1A^Y235R^, STX1A^E238V^, STX1A^V248K^ and STX1A^S249E^) and double point mutations (STX1A^D231N,R232N^ and STX1A^V248K,S249E^) if the residues were in sequential positions and analyzed their expression at the synapse and the corresponding levels of Munc18-1 at the synapse (Figure 6A, Figure 6-figure supplement 1). Notably, we excluded STX1A^A240S^, which is present in STX1B, redundant in function to STX1A (Vardar et al., 2016) and STX1A^R246^ and STX1A^D250^ which are electrochemically similar to STX2^H246^ and STX2^E250^ (Figure 6A). For unknown reasons, the levels of some of the mutant constructs of STX1A seem to be reduced more than 50%, which has been shown to be detrimental for release (Arancillo et al., 2013; Vardar et al., 2016) while all electrophysiological parameters showed comparable levels to STX1A WT or even an enhanced spontaneous release (Figure 6). For this reason, we did not explore the expression results any further. Additionally, levels in Munc18-1 of the corresponding mutants were also decreased (Figure 6-figure supplement 1).

**Figure 6.**
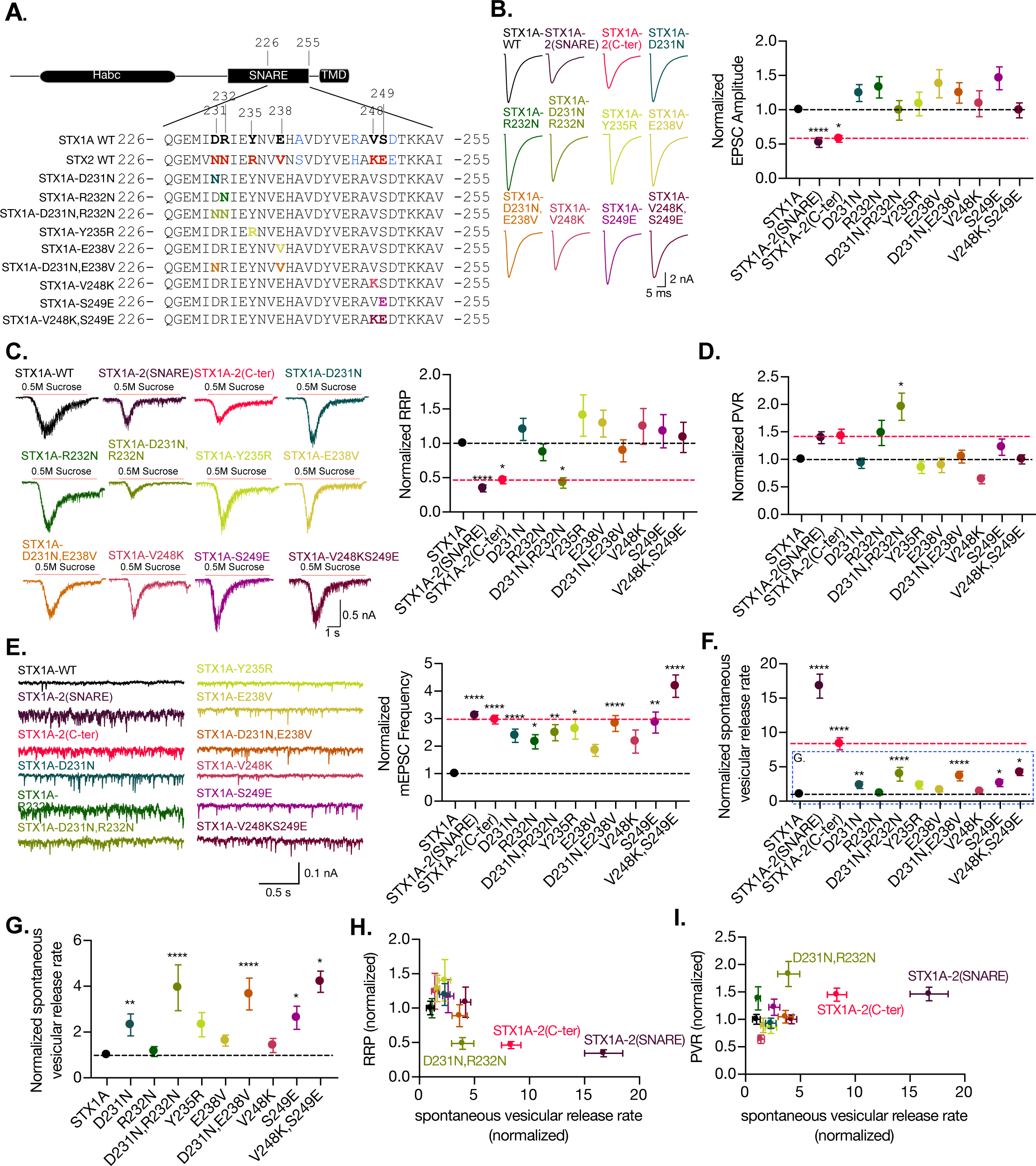
The charge of the outer-surface residues in the C-terminal half of the SNARE domain is important for clamping spontaneous release and D231,R232 are important in the stabilization of the pool and the efficiency of Ca^2+^-evoked release. A. Sequence of the C-terminal half of STX1A and STX2 and single- and double-point mutations in the sequence of STX1A WT. B. Example traces (left) and quantification of the EPSC amplitude (right) from autaptic STX1A-null hippocampal mouse neurons rescued with STX1A, STX1A^D231N^, STX1A^R232N^, STX1A^Y235R^, _STX1A_E238V_, STX1A_V248K_, STX1A_S249E _STX1A_D231N,R232N _and STX1A_V248K,S249E. C. Example traces (left) and quantification of the response induced by a 5s 0.5M application of sucrose, which represents the readily releasable pool of vesicles (RRP). D. Quantification of the vesicle release probability (PVR) as the ratio of the EPSC charge over the RRP charge (PVR). E. Example traces (left) and quantification of the frequency of the miniature excitatory postsynaptic currents (mEPSC) (right). F. Quantification of the spontaneous vesicular release rate as the ratio between the of mEPSC frequency and number of vesicles in the RRP. G. Quantification of the spontaneous vesicular release rate as the ratio between the of mEPSC frequency and number of vesicles in the RRP but STX1A-2(SNARE) and STX1A-2(Cter) chimeras were removed. H. Correlation between the RRP and the spontaneous vesicular release rate. Both values are normalized to STX1A WT. I. Correlation between the PVR and the spontaneous vesicular release rate. Values are normalized to STX1A WT. Data information: (B-H) All electrophysiological recording were done on autaptic neurons. Between 30-35 neurons per group from 3 independent cultures where recorded. Values from STX1A-2(SNARE) and STX1A-2(Cter) groups were taken from the experiments done in Figure 2 and normalized to their own STX1A WT control and used here for visual comparison. Data points represent the mean ±SEM. Significances and P values of data were determined by non-parametric Kruskal-Wallis test followed by Dunn’s post hoc test; *p≤0.05, **p≤0.01, ***p≤0.001, ****p≤0.0001. All data values are summarized in Figure 6 – Source Data 1.

The electrophysiological analysis of our point mutations showed that none of the mutations caused a change in the EPSC amplitude (Figure 6B). However, the double point mutant STX1A^D231N,R232N^ was the only mutant that showed a decrease in the RRP size from 0.469nC (SEM±0.08) to 0.19nC (SEM±0.03) (Figure 6C), as well as a 2-fold increase in the PVR (Figure 6D). This is consistent with the changes in our previous findings from the STX1A chimeric analysis, which showed a decreased RRP and an increased PVR (Figure 2G). All mutants showed an increase in the mEPSC frequency; however, it was not significant for STX1A^E238V^ and STX1A^V248K^ (Figure 6E). This suggests that any small alteration in the outer surface of the C-terminal half of the SNARE domain of STX1A could de-stabilize the clamp for spontaneous release. Therefore, the whole domain is crucial in the regulation of the synaptic vesicle clamp. The mutants with increased spontaneous release frequency also exhibited an increase in the release rate (Figure 6F and G), particularly those with a simultaneous increase in the mEPSC frequency and a trend towards RRP reduction. This included all the double point mutants STX1A^D231N,R232N^, STX1A^D231D,E238V^ and STX1A^V248K,S249E^ and the single point mutants STX1A^D231^ and STX1A^S249E^. These results suggest that the more alterations the C-terminal half of the SNARE domain undergoes, the less effective it is in maintaining the stability of the vesicles in the RRP and in clamping spontaneous release.

Finally, we examined the relationship between RRP size and release rate (Figure 6I) and PVR and release rate (Figure 6H). Changes that de-stabilize the primed state might be seen as a decrease in the RRP and an increase the release rate. Changes in the height of the energy barrier for fusion may make it harder or easier for vesicles to fuse and could be indicated by a decrease or increase in the PVR, respectively, accompanied by a change in the release rate. Surprisingly, STX1A-2(SNARE), STX1A-2(Cter) and STX1A^D231N,R232N^ exhibit changes in both these correlations: decreased RRP and increased spontaneous release rate (Figure 6I), and increased PVR and spontaneous release rate (Figure 6H). This may suggest D231, R232 may have a role on the stability of primed vesicles and on the energy barrier for fusion. However, the integrity of the SNARE domain of STX1A is more important than single-point changes, suggesting an additive effect of the residues of the C-terminal half of the SNARE domain.

## Discussion

In this study we investigated the role and specificity of the SNARE domain of STX1 in regulating neurotransmitter release. We found that the C-terminus of the SNARE domain of STX1 is crucial in the regulation of multiple aspects of synaptic transmission, such as the clamping of spontaneous release and the formation and maintenance of the RRP of vesicles. Furthermore, it is involved in the regulation of speed and efficacy of Ca^2+^-evoked release, however, these functions are dependent on the integrity of the full-SNARE domain of STX1 and regions outside of the SNARE domain.

### STX1 is fine-tuned for synaptic vesicle release

Current understanding of constitutive and regulated membrane fusion involves the idea of the energetically favorable interaction of four cognate SNARE domains that “zipper-up” and pull the vesicle and plasma membrane together (Fasshauer et al., 1998). In the physiological context, vesicle fusion can vary dramatically in terms of efficacy and acceleration upon triggering, arguing for an extensive regulation machinery for the fusion process. An important role is attributed to regulatory proteins, such as complexins and synaptotagmins, that regulate various aspects of vesicle fusion including synaptic vesicle docking and priming, clamping of spontaneous release and speed and efficacy of neurotransmitter release (Rizo, 2022; Rizo & Rosenmund, 2008; Südhof, 2013). Although SNARE proteins are quite promiscuous in their assembly (Bajohrs et al., 2005; Brunger, 2005; Fasshauer et al., 1998; Peng et al., 2013; Vardar et al., 2022), variations in their SNARE motif sequence can lead to differential regulation of the fusion reaction beyond their role in zippering, e.g. by modifying the surface charge (Kádková et al., 2023; Ruiter et al., 2019) or regulating interaction with modulatory proteins (Schupp et al., 2016; Stepien et al., 2022; Zhou et al., 2015). We show that STX2 can rescue some aspects of neurotransmitter release in STX1-null neurons, supporting redundant functions among syntaxin isoforms in executing vesicle fusion. However, regulated fusion with non-cognate trans-SNARE complexes containing STX2 showed several unfavorable release properties, including slowed and inefficient Ca^2+^-evoked release, a reduced pool of fusion competent vesicles, while spontaneous release was greatly increased (Figure 1). Given these findings, and our previous findings which compared the release parameters of the tonic release isoform STX3 with STX1 (Vardar et al., 2022), we argue that STX1 is finetuned for phasic release with a high signal-to-noise ratio for evoked over spontaneous release.

### The C-terminus of the STX1 SNARE domain is important for stability of the primed state of vesicles and the clamping of spontaneous fusion

As a general observation, the exchange of the whole or only the C-terminal half of the SNARE domains of STX1 and STX2 resulted in altered Ca^2+^-dependent responses, PVR, PPR, short-term depression, as well as release rate (Figure 2 and 3). It is clear that the C-terminus of the STX1 SNARE domain is important for the stability of the primed state of the vesicles and the clamping of spontaneous fusion. While conducting a sub-saturating sucrose experiment (Basu et al., 2007; Ruiter et al., 2019; Schotten et al., 2015) could further confirm the hypothesis of reduced fusogenicity in STX1 chimeras, the small RRP obtained at 500 mM sucrose poses a limitation to our study. Therefore, we must rely on other parameters to interpret these results. Generally, changes in the molecular mechanisms for priming proportionally change the Ca^2+^-evoked response while leaving the PVR unchanged. Changes in the PVR, PPR, release rate and depression during high-frequency train stimulation, however, may indicate alterations in the fusogenicity of the vesicles that stem from aberrations in the mechanisms underlying the regulation of the fusion energy barrier (Schotten et al., 2015). In this light, we propose a model where the C-terminal half of the SNARE domain of STX1 plays a major role in the stabilization of the primed state and in the clamping of the spontaneous release. Disrupting the C-terminus of the STX1A SNARE domain by adding that of the non-cognate STX2 (Figure 7-red dotted line) may impact vesicle fusogenicity by destabilizing the primed state and decreasing the fusion energy barrier that now becomes easier for vesicles to overcome (Schotten et al., 2015). Additionally, adding the C-terminus of the SNARE domain of STX1A to STX2 may increase the ability of the SNARE complex to stabilize the primed state by increasing the degree of the fusion energy barrier and thereby establish a clamp for spontaneous release (Figure 7-light green dotted line). This model supports our previous findings in which, by performing a similar chimeric analysis with STX3, we suggested the C-terminus of the SNARE domain of STX1 as a key regulator in the clamping of spontaneous vesicle fusion (Vardar et al., 2022). We discuss 3 possible mechanisms through which the C-terminus of the SNARE domain of STX1 might contribute to the stability of the primed state of synaptic vesicles and the clamping of spontaneous release.

**Figure 7.**
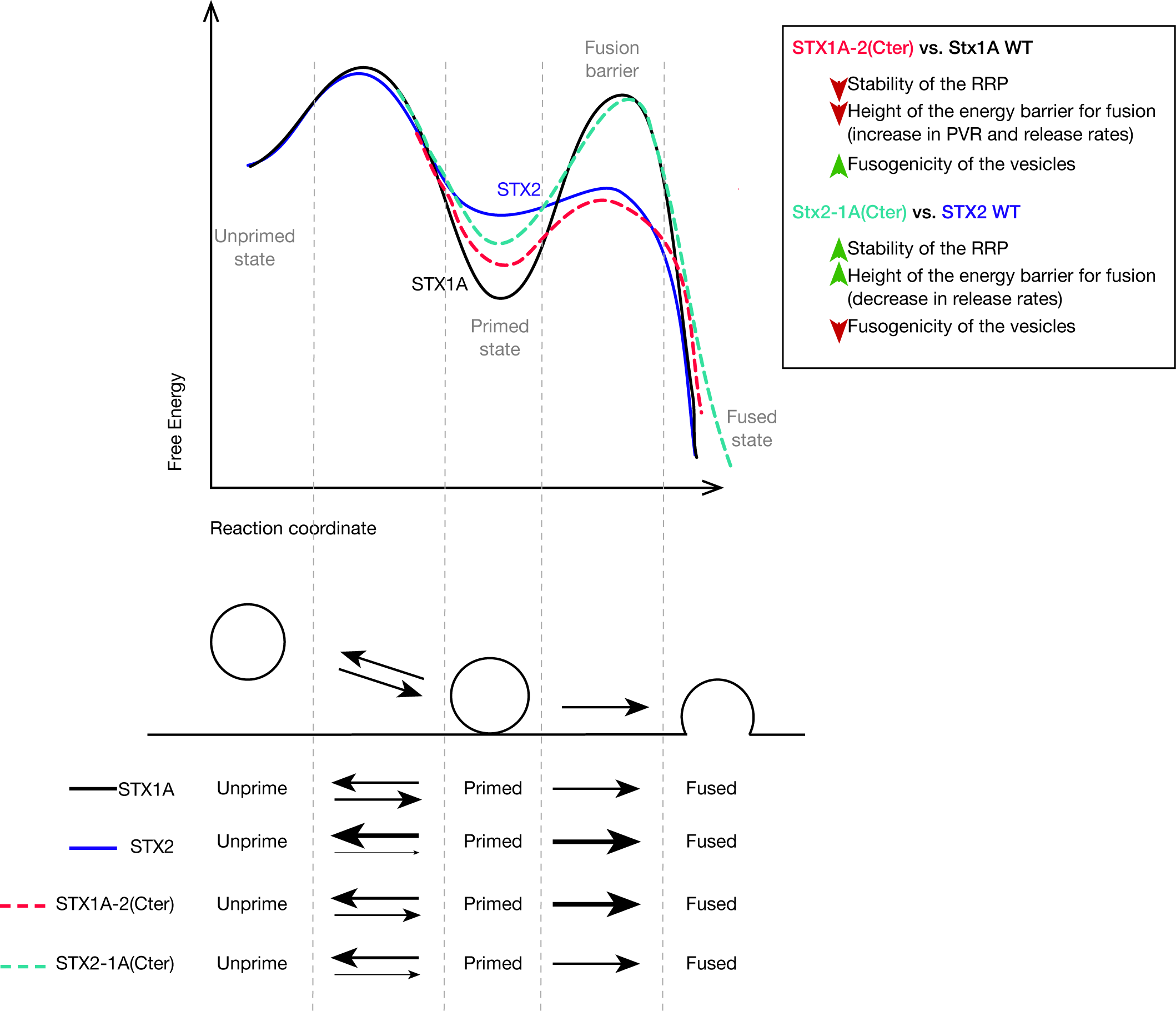
Speculative model based on most important changes in electrophysiological properties found in STX1A-chimeras or STX2-chimeras compared to their WT isoforms. A. Energy landscape for priming and fusion of synaptic vesicles that have SNARE complexes formed with STX1A WT (black line), STX1A-2(SNARE or Cter) (red dotted line), STX2 WT (blue line); STX2-1A(Cter) (light green dotted line). B. Summarized conclusions of our results based on the speculative model in (A). C. Changes in the equilibrium between the unprimed/primed state (reflecting the stability of the primed state) and the rate of fusion (energy landscape for fusion adapted from “Sørensen, 2009”).

### The interaction of the C-terminus of the SNARE domain of STX1A with Munc18-1 in the stabilization of the primed pool of vesicles

Previous studies have attributed the vesicle priming function to the N-terminal half of the SNARE domain (Sørensen et al., 2006; Weber et al., 2010) and to proteins involved in the N-terminal nucleation of the SNARE complex, such as Munc13 and Munc18-1 (Ma et al., 2011; Wang et al., 2020). One line of thought proposes that these proteins prime vesicles to the plasma membrane by forming a SYB2-STX1 acceptor-complex for SNAP-25 (Jiao et al., 2018; Lai et al., 2017; Parisotto et al., 2012; Stepien et al., 2022; Wang et al., 2019). Thus, changes in Munc18-1 levels in STX1 and STX2 chimeras is one of the first hypotheses that comes to mind as an underlying mechanism of the altered primed state of the vesicles. Notably, we did not observe changes in the levels of Munc18-1 when exchanging N-terminal halves of the SNARE domain between syntaxins. However, all the chimeras that contained the C-terminal half of the SNARE domain of the non-cognate isoform showed an increase in levels of Munc18-1 compared to their native form (Figure 4C, Figure 5F) while manifesting opposing effects in the stabilization of primed vesicles: STX2 chimeras showed a better stabilized primed state reflected in the increase of the RRP and a decrease in the fusogenicity parameters (Figure 5H-J and Figure 7), whereas STX1A-chimeras showed signs of a destabilized primed state (Figure 4F-H and Figure 7). These opposing phenotypes both accompanied by an increase in Munc18-1 levels might be the manifestation of two modes of binding of Munc18-1 to the C-terminal half of the SNARE domain of STX1. First, Munc18-1 was shown to maintain an interaction with the C-terminus of the SNARE domain of STX1 during the N-terminal nucleation of the SNARE complex, important for the initiation of the SNARE complex assembly (Stepien et al., 2022). Additionally, the central cavity of Munc18-1 interacts with the C-terminal half of the SNARE domain of STX1 in its closed conformation, important for the chaperoning and trafficking of STX1 to the plasma membrane (Han et al., 2011, Shi et al., 2021). Mutations of the corresponding residues in STX1 (including D231 and E234), while affecting the binding and chaperoning functions of Munc18-1, did not affect the fusion in liposome fusion assays (Shi et al., 2021). We can speculate that the presence of the C-terminal half of the SNARE domain of STX2 might favor an open conformation of syntaxin due to an inability of Munc18-1 to chaperone the closed conformation and the open conformation of STX1 was previously shown to display an increase in fusogenocity parameters (Gerber et al., 2008; Zhou et al., 2013; Vardar et al., 2021). On the other hand, the presence of the C-terminus half of the SNARE domain of STX1 allows for the binding of Munc18-1 and favors the chaperoning of a closed conformation of syntaxin, which would display reduced fusogenicity parameters. Therefore, it is tempting to speculate that Munc18-1 might have a qualitative effect on the vesicle stabilization rather than a quantitative one. In this case, it can be argued that an intact C-terminal half of the SNARE domain is a conditional requirement in order for Munc18-1 to fulfill ist chaperoning and nucleation function on the N-terminus in a physiological setting, thus leading to a reduced RRP (Figure 4F).

### The participation of STX1A in the electrostatic regulation of the fusion energy barrier model

Besides the destabilization of the RRP, we found that the clamping of the spontaneous release is particularly susceptible to changes in single or double consecutive residues in the C-terminus of the SNARE domain (Figure 6). The degree of the vesicle fusion energy barrier is largely regulated by the electrostatic landscape of the fusion apparatus (Shao et al., 1997; Trimbuch et al., 2014; Williams et al., 2009) which will try to neutralize the negatively charged approaching membranes so they can fuse. This is also influenced by the total charge of the SNARE domain complex (Ruiter et al., 2019). Thus, changes in the net charge of the SNARE complex can cause altered vesicle-fusogenicity and as a result, altered release rates at the synapse (Ruiter et al., 2019). The observation of SNAP-25 charge-reversal mutations proposed a model by which a positive net charge increase lowerers the fusion energy barrier and increases fusion rates because it neutralizes the negative charges of the membranes, while a negative net charge increase increases the negative electrostatic fusion energy barrier causing a decrease in fusion rates (Kádková et al., 2023; Ruiter et al., 2019). In this light, it is plausible to argue that the differences in the release behavior led by the single or double-point mutations in the C-terminal half of the SNARE domain of STX1A (Figure 6) might be due to the altered electrostatic net charge. The net charge difference in the C-terminal half of the SNARE domain between STX1A and STX2 is only one positive charge that would only lead to a minor change in the release rate. However, the release rate we observed are 15-fold (in the native form) (Figure 1K) or 17-fold and 8-fold (in STX1A chimeras) (Figure 2H) increased. Additionally, the mutations STX1A^D231N,R232N^ and Stx1A^V248K,S249E^ which result in a null net charge difference, increased the spontaneous release rate 4-fold compared to STX1A WT, or mutation STX1A^S249E^ that should reduce the release rate according to this hypothesis as it possesses one negative net charge difference increases the spontaneous release rate 3-fold (Figure 6G). Therefore, our results suggest that the changes in release rate in STX1A point-mutations are not caused merely by electrostatic changes but by an independent mechanism from the electrostatic model of regulation of releas.

### The interaction of STX1A with the C2B domain of SYT1 through the primary interface

What can be the mechanism through which the C-terminal half of the SNARE domain regulates vesicle fusion if it is not solely through electrostatic changes? Interestingly, neurons expressing STX2 showed a slowed down Ca^2+^-evoked release, a dramatic increase in the spontaneous release rate, and decrease in the RRP (Figure1). Additionally, STX1A-chimeras which contained the C-terminus of the SNARE domain of STX2 showed a decrease in the RRP and a dramatic unclamping of the spontaneous release rate (Figure 2). This is reminiscent of the well-established phenotype of SYT1-null mouse neurons (Bouazza-Arostegui et al, 2022; Stepien et al., 2022; Xu et al, 2007; Xue et al., 2009) and suggests a possible alteration in the function of SYT1 in these neurons. Moreover, mutating individual residues in the C-terminal half of the SNARE domain which differ between STX1A and STX2 showed a particular effect on the clamping of spontaneous release and the RRP in the case of D231 and R232 (Figure 6E), two functions associated to SYT1-SNARE interaction (Schupp et al., 2016). Some of these residues (D231 and E238) have been identified by crystal structure studies to be involved in the putative interaction between the C2B domain of SYT1 and the C-terminal half of the SNARE domain of STX1 and SNAP-25, referred to as the primary interface (Zhou et al., 2015). Mutations of the residues at the primary interface in SYT1 and SNAP-25 identified this putative interaction as a key regulation site for the clamping of spontaneous release and Ca^2+^-triggered release (Chang et al., 2018; Schupp et al., 2016; Zhou et al., 2015). The partial rescue of the speed of Ca^2+^-evoked release, the RRP and the clamping of the spontaneous release in STX2 chimeras which contained the C-terminus of the SNARE domain of STX1A (Figure 3), suggested a gain of function though the interaction with SYT1. However, none of the changes in the C-terminus of the SNARE domain of STX1A resulted in a change in the speed of release (Figure 2 and Figure 6) and the release rates obtained in STX1A-2(SNARE) and STX1A-2(Cter) were dramatically higher (Figure 2) than in SYT1-KO neurons (Bouazza-Arostegui et al., 2022). In this light, our data support the primary interface as a key site for the regulation of spontaneous release, in addition to further unidentified mechanisms involving other residues in the C-terminal half of the SNARE domain of STX1A. Notably, it argues against STX1 as part of the Ca^2+^-evoked release mechanisms through its interaction with the C2B domain of SYT1, supporting our previous findings that identified regions outside of the SNARE domain of STX1, such as the juxtamembrane and transmembrane domain, as regulators of the synchronization of neurotransmitter release (Vardar et al., 2022).

As a final remark, it is possible that the changes in the spontaneous release rate and the priming stability may stem from a reduced stability of the SNARE complex itself through putative interactions between outer surface residues. Studies of the kinetics of assembly of the SNARE complex which mutate solvent-accessible residues in the C-terminal half of the SNARE domain of SYB2 have shown reduction in the stability of the SNARE complex assembly and are correlated with impaired fusion (Jiao et al., 2018). However, STX1 mutations of outward residues were inconclusive and were always accompanied by hydrophobic layer mutations (Jiao et al., 2018), which affect the assembly kinetics and energetics of the SNARE complex (Ma et al., 2015). Single molecule optical-tweezer studies have focused on the impact of regulatory molecules on the stability of assembly such as Munc18-1 (Ma et al., 2015; Jiao et al., 2018) and complexin (Hao et al., 2023), or on the intrinsic stability of the hydrophobic layers in the step-wise assembly of the SNARE complex (Gao et al., 2012; Ma et al., 2015; Zhang et al., 2017). Although the conserved hydrophobic layers in the SNARE domains of STX1A and STX2 (Figure 1) suggest unchanged zippering and intrinsic stability of the complex, further studies addressing the contribution of surface residues on the stability of the alpha-helical structure of the SNARE domain of STX1 (Li et al., 2022) or the stability of the SNARE complex should be conducted.

## Materials and methods

### Animal maintenance and generation of mouse lines

All animal-related procedures and experiments were performed in accordance with the guidelines and approved by the animal welfare committee of Charité-Universitätsmedizin and the Berlin state government Agency for Health and Social Services, under license number T0220/09. To generate the STX1-null mouse model, we bred the conventional *Stx1A*-knockout (KO) line, in which exon 2 and 3 were deleted (Gerber et al., 2008), with the *Stx1B* conditional-KO line in which exons 2–4 were flanked by loxP sites (Wu et al., 2015).

### Neuronal cultures

Primary hippocampal neuronal cultures were prepared from STX1A/1B cDKO mice at postnatal day 0-1 and were seeded onto a continental astrocyte feeder layer (for immunocytochemistry 40K neurons/well in 12-well plates) or astrocyte micro-dot islands (for the electrophysiology 4K neurons/well in 6-well plates) as previously described (Vardar et al., 2016; Xue et al., 2007). Cortical neuronal cultures were prepared from the same mice and plated onto continental astrocyte feeder layer for Western Blot experiments (500K neurons/ well in 6 well plates). Cultures were incubated in NeuroBasal-A (NBA) medium (Invitrogen) supplemented with B-27 (invitrogen), 50 IU/ml penicillin and 50μg/ml streptomycin) at 37°C before and during the experiments. Neuronal cultures were incubated for 13-20 days *in vitro* (DIV).

### Lentiviral construct production and viral infection

Point mutations were introduced into *Stx1A* or *Stx2* using a TA cloning vector with the QuickChange Site-Directed Mutagenesis Kit (Stratagene). Chimeric constructs of both syntaxin isoforms were obtained using Gibson Assembly (NEB). For the viral infection the cDNA of mouse *Stx1A* (NM_016801.4), *Stx2* (NM_007941.2), chimeric and point-mutation constructs were cloned into a lentiviral shuttle vector containing a nuclear localization signal (NLS) GFP-P2A expression cassette under the human synapsin-1 promoter. For obtaining the *Stx1B*-KO of in the *Stx1A*(*-/-*);*Stx1B*(*flox/flox*) neurons (Stx1-null), we used a lentiviral construct which induces the expression of improved Cre-recombinase (iCre) fused to NLS-RFP-P2A under the control of human synapsin-1 promoter. All viral particles were made by the Viral Core Facility of Charité-Universitätsmedizin as previously described (Lois et al., 2002). Empty NLS-GFP-P2A constructs were used as control. For all experiments cultures were infected with lentiviral particles at DIV1-2.

### Neuron viability

For neuron survival assays we quantified the number of the STX1-null hippocampal neurons transduced with *Stx1A, Stx2 and GFP* (as control) at DIV 15, 22, and 29, and compared this to the number of neurons at DIV8. Two wells per group were plated and fifteen random ROIs of 1.23mm2 per well were imaged at the different time points (thirty total ROIs per group at each time point per culture). Phase-contrast bright field images and fluorescent images with excitatory lengths of 488 and 555 nm were acquired on a DMI 400 Leica microscope, DFC 345 FX camera (Leica), HCX PL FlUPTAR 10x objectives (Leica) and LASAD soft-ware (Leica). The neurons were counted offline witht the 3D Object Counter function in Fiji software (Vardar et al., 2016). For the example images the cultures were fixed at the corresponding time points in 4% paraformaldehyde (PFA) in 0.1M phosphate-buffered saline, pH 7.4 (PBS) and immunocytochemistry was done as follows.

### Immunocytochemistry and image acquisition

High-density hippocampal neuron cultures (40 x 103) were plated onto the astrocyte-feeder layer and infected at DIV1-2. At DIV14-15 the cultures were fixed in 4% PFA in PBS for 10 min, permeabilized in 0.1% Tween-20 PBS (PBS-T) for 45 min at room temperature (RT) and blocked in 5% normal donkey serum in PBS-T for 1h at RT. To detect our proteins of interest the neurons are incubated with the primary antibodies in PBS-T at 4°C over-night (O/N). For protein expression analysis neurons were incubated with guinea-pig polyclonal anti-VGlut1(1:4000; Synaptic Systems), mouse monoclonal anti-STX1A (1:1000; Synaptic Systems) and rabbit polyclonal anti-STX2 (1:1000; Synaptic Systems) or rabbit polyclonal Munc18-1 (1:1000; Sigma-Aldrich). Secondary antibodies conjugated with rhodamine red, or Alexa Fluor 488, or 647 (1:500; Jackson ImmunoResearch) in PBS-T were applied for 1h at RT in the dark. For the example images for survival assay neurons were incubated with chicken polyclonal anti-MAP2 (1:2000; Merck Milipore) and secondary antibody Alexa Fluor 488. The coverslips were mounted on glass slides with Mowiol mounting agent (Sigma-Aldrich). For quantitative analysis, every comparable group in each culture is treated with the same antibody solution. Images are acquired with an Olympus IX81 epifluorescence-microscope with MicroMax 1300YHS camera (Princeton Instruments) and with MetaMorph software (Molecular Devices). Images for the example images of the survival assay and for the quantitative analysis of protein expression were taken with an optical magnification of 10X and 60X, respectively. The exposure times of each wavelength are kept constant for all experimental groups in one culture. Overexposure and photobleaching is avoided by monitoring the fluorescent saturation level at the synapse. To image glutamatergic neurons 7-9 images that had a fluorescent signal for VGlut1 staining are taken from each group in each experimental replicate. Data was analyzed offline on ImageJ. Three to five regions per image were selected which showed neuronal projections marked by VGlut1-positive labelling. Vglut1-positive puncta are selected using MaxEntropy threshold function of the selected regions, creating a VGlut1-positive ROI. The intensities of VGlut1, STX1A, STX2 and Munc18-1 are measured using the corresponding ROIs. For the quantification of relative protein expression at glutamatergic synapses, the ratio between STX1A, STX2 or Munc18-1 to VGlut1 is calculated in each selected region. To compare between groups the data is normalized to STX1A group or STX2 group.

### Electrophysiology

To assess synaptic function, whole-cell patch-clamp recordings in autaptic neurons were performed at DIV13-DIV19 at room temperature. Synaptic currents were recorded using a MultiClamp 700B amplifier (Molecular Devices) and data was digitally sampled at 10kHz and low-pass filtered at 3kHz with an Axon Digidata 1440A digitizer (Molecular Devices). Data acquisition was controlled by Clampex10 software (Molecular Devices). Series resistance was compensated at 70% and only neurons with a series resistance lower than 10MΩ were further recorded. Neurons are continuously perfused using a fast perfusion system (1-2ml/min) with an extracellular solution (unless stated otherwise) that contains 140mM NaCl, 2.4mM KCl, 10mM HEPES, 10mM glucose, 2mM CaCl2 and 4mM MgCl (295-305mOsm, pH7.4). Borosilicate glass pipettes are pulled yielding a resistance between 2-5MΩ. Pipettes are filled with a KCl-based intracellular solution containing 136mM KCl, 17.8mM HEPES, 1mM EGTA, 4.6mM MgCl2, 4mM Na2ATP, 0.3mM Na2GTP, 12mM creatine phosphate, and 50 Uml1 phosphocreatine kinase (300mOsm; pH7.4). Only neurons that express RFP and GFP are selected for recording and cells with a leak current higher than 300pA were excluded. Neurons are clamped at −70mV during all protocols of electrophysiological recording. For triggering action potential (AP) evoked release neurons are stimulated by a 2ms depolarization to 0mV and excitatory postsynaptic currents (EPSCs) are registered. To measure the synchronicity and the kinetics of the AP-evoked responses we inverted the EPSC change and integrated our signal For measuring spontaneous release (mEPSC) we recorded at −70mV for 48s and for 24s in the presence of the AMPA-receptor antagonist NBQX (3μM) (Tocris Bioscience) diluted in extracellular solution. To calculate the frequency of mEPSC we filtered at 1kHz and analyzed using a template-based miniature event detection algorithm implemented in the AxoGraph X software. We then subtracted the measured mEPSC in NBQX to the mEPSCs measured in extracellular solution. The release of the readily releasable pool (RRP) of synaptic vesicles is triggered by the application of a 500mM hypertonic sucrose solution diluted in extracellular solution for 5s (Rosenmund and Stevens, 1996), and the RRP size is estimated by integrating the area of the sucrose-evoked current setting as baseline the steady-state current. The vesicular release probability (PVR) of each neuron is determined by the ratio between the charge of the EPSC and the size of the RRP. The spontaneous release rate is calculated as the ratio between the mEPSC frequency and the number of primed vesicles. This determines the fraction of the RRP which is spontaneously released per second. To measure the cumulative charge transfer of the synaptic responses we inverted the EPSC charge and integrated the signal. Short term plasticity was examined either by evoking 2 AP with 25 ms interval (40 Hz) or a train of 50 AP at an interval of 100 ms (10 Hz). Data were analyzed offline using Axograph10 (Axograph Scientific)

### Western Blot

Protein lysates from cortical mass cultures were prepared by lysing neurons in 200 μl lysis buffer containing 50 mm Tris/HCl, pH 7.9, 150 mm NaCl, 5 mm EDTA, 1% Triton X-100, 0.5% sodium deoxycholate, 1% Nonidet P-40, and 1 tablet of Complete Protease Inhibitor (Roche) for 30 min on ice. Samples are boiled for 5min at 95°C. Equal amounts of protein were loaded onto a 12% SDS-PAGE and run in electrophoresis buffer at 80V for 30 min and then 120mV for 1h. Proteins were transferred onto a nitrocellulose membrane at 50mA O/N. The membranes were blocked in 5% milk for 1h at room temperature and incubated with the corresponding primary antibody diluted in PBS for 1h at room temperature. Membranes were incubated with mouse monoclonal anti-STX1A (1:10,000; Synaptic Systems), rabbit polyclonal anti-STX2 (1:10,000; Synaptic Systems), rabbit polyclonal anti-STX3A (1:10,000; Synaptic Systems), rabbit polyclonal anti-Munc18-1 (1:10,000; Sigma-Aldrich) and mouse monoclonal anti-betaTubIII (1:10,000; Sigma) (as loading control). Secondary antibodies conjugated with HRP-conjugated goat secondary antibodies (1:10,000; Jackson ImmunoResearch) diluted in PBT were applied for 1h at RT. Membranes are then incubated with ECL Plus Western Blotting Detection Reagents (GE Healthcare Biosci-ences) and luminal signal was visualized and imaged in Fusion FX7 image and analytics system (Vilber Lourmat).

### Statistical analysis

Data presented in box-whisker plots are compiled as single observations, median, quartiles and outliers. Data in bar graphs and X-Y plots present means ± SEM. All data were tested for normality with D’Agostino-Pearson test. Data from two groups with non-parametric distribution were subjected to Mann-Whitney test. Data from two groups with normal distribution were subjected to unpaired two-tailed t-test. Data from more than two groups were subjected to Kruskal-Wallis followed by Dunn’s *post hoc* test when at least one group showed a non-parametric distribution. Data from more than two groups were subjected to Ordinary one-way ANOVA when all groups showed a parametric distribution. STP was subjected to two-way ANOVA. Statistical analyses were performed using Prism 7 (GraphPad). All statistical data are summarized in the corresponding “Source Data” tables.

Fold-increase/decrease and percentage-increase/decrease were done using the mean values of each parameter.

## Acknowledgements

We thank the Charité Viral Core facility, Katja Pötschke, and Bettina Brokowski for virus production, Berit Söhl-Kielczynski and Heike Lerch for technical assistance, Melissa Herman for their contribution to the manuscript and to all the Rosenmund Lab members for the discussions. This project was funded by the German Research Councin (DFG) grants 399894546, 184695641, 278001972 and 390688087 and by a SFB958 doctoral fellowship.

---

**Figure 1 – figure supplement 1.**
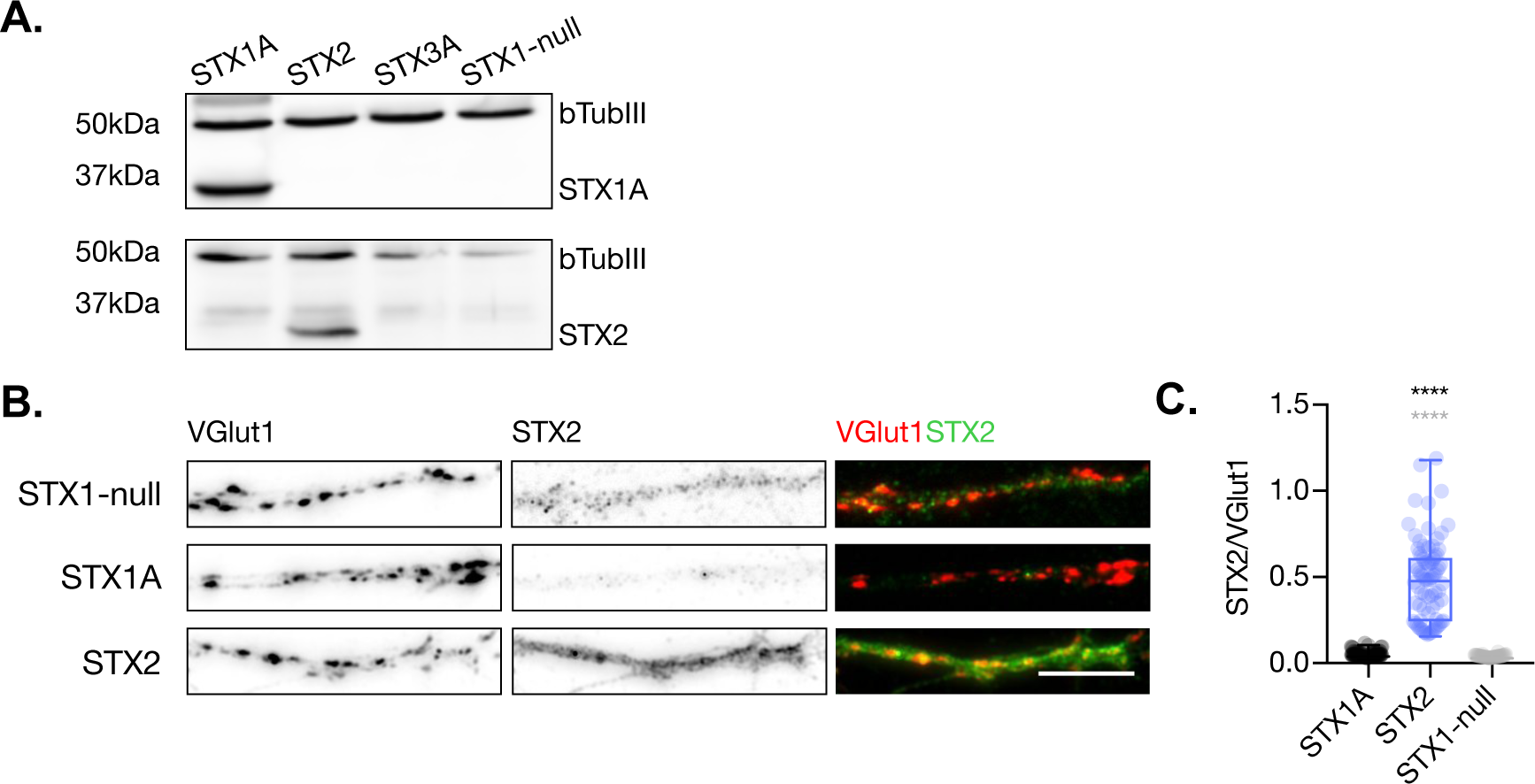
STX2 is expressed in STX1-null hippocampal neurons. A. SDS-PAGE electrophoresis of lysates from STX1-null neurons infected with STX1A, STX2 or GFP (STX1-null) and STX3A as negative controls. Proteins were detected using antibodies that recognize β-Tubuline III as loading control STX1A, STX2 and STX3A. B. Example images of mass-culture hippocampal neurons stained with fluorophore-labeled antibodies that recognize VGlut1 (red in merge) and STX2 (green in merge) from left to right. Scale bar: 10μm. C. Quantification of the immunofluorescent intensity of STX2 normalized to the intensity of the same VGlut1-labeled ROIs. Data information: In (C) data is show in a whisker-box plot. Each data point represents single ROIs, middle line represents the median, boxes represent the distribution of the data, where the majority of the data points lie and external data points represent outliers. Significances and P values of data were determined by non-parametric Kruskal-Wallis test followed by Dunn’s post hoc test; *p≤0.05, **p≤0.01, ***p≤0.001, ****p≤0.0001. All data values are summarized in Figure 1 – Figure Supplement 1 – Source Data 1.

**Figure 2 – figure supplement 1.**
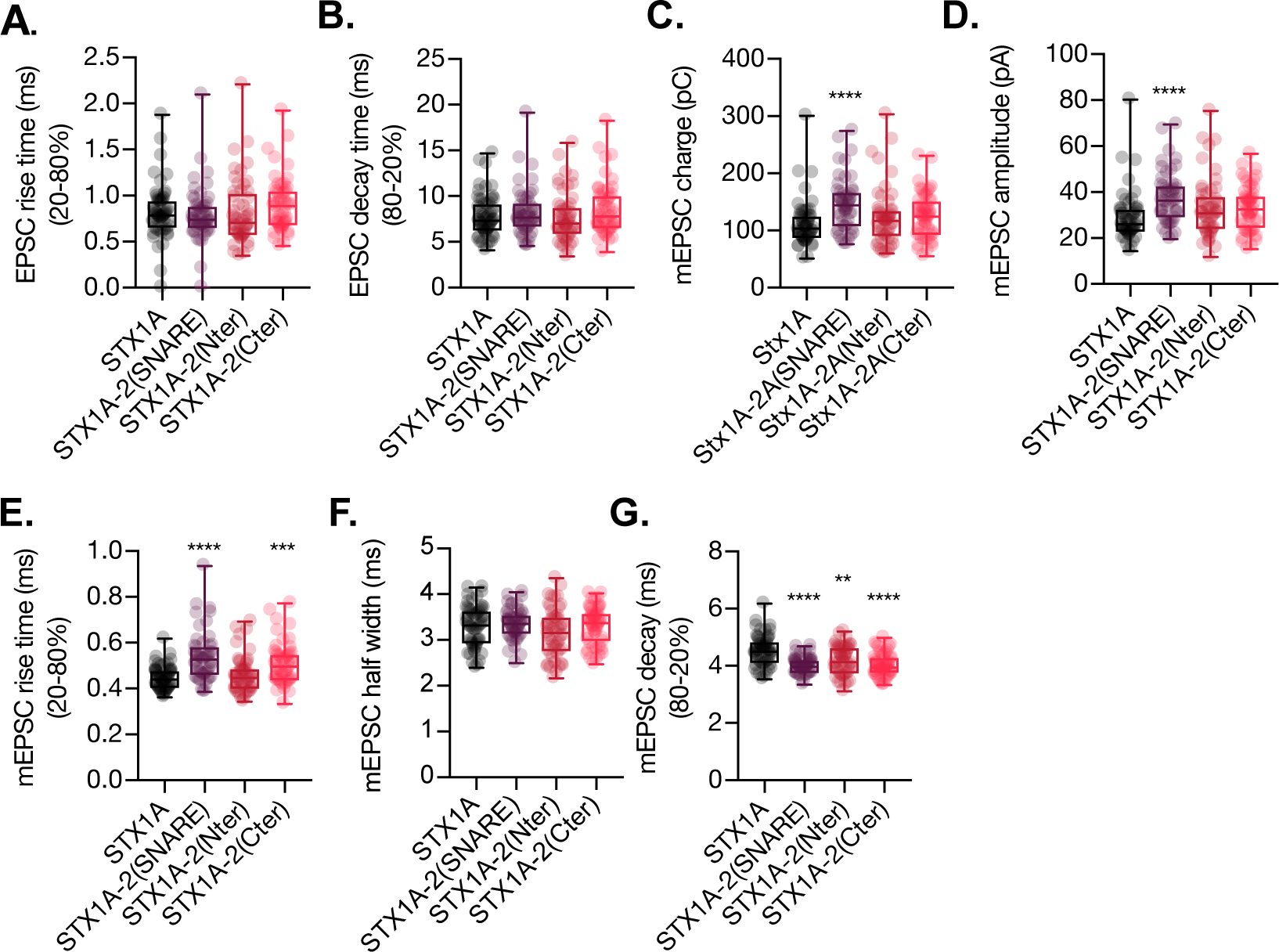
Quantification of kinetic parameters of the EPSC and the mEPSC of autaptic STX1A-null hippocampal mouse neurons rescued with STX1A, STX1A-2(SNARE), STX1A-2(Nter) or STX1A-2(Cter). A. Quantification of the rise time (20-80%) of the EPSC neurons B. Quantification of the decay time (80-20%) of the EPSC. C. Quantification of the mEPSC charge. D. Quantification of the mEPSC amplitude. E. Quantification of the rise time (20-80%) of the mEPSC. F. Quantification of the half width of the mEPSC. G. Quantification of the decay time of the mEPSC. H. All electrophysiological recording were done on autaptic neurons. Data information: Data in (A-G) is show in a whisker-box plot. Each data point represents single observations, middle line represents the median, boxes represent the distribution of the data, where the majority of the data points lie and external data points represent outliers. Significances and P values of data were determined by non-parametric Kruskal-Wallis test followed by Dunn’s post hoc test or by Ordinary one-way ANOVA; *p≤0.05, **p≤0.01, ***p≤0.001, ****p≤0.0001. All data values are summarized in Figure 2 – figure Supplement 1 – Source Data 1

**Figure 3 – figure supplement 1.**
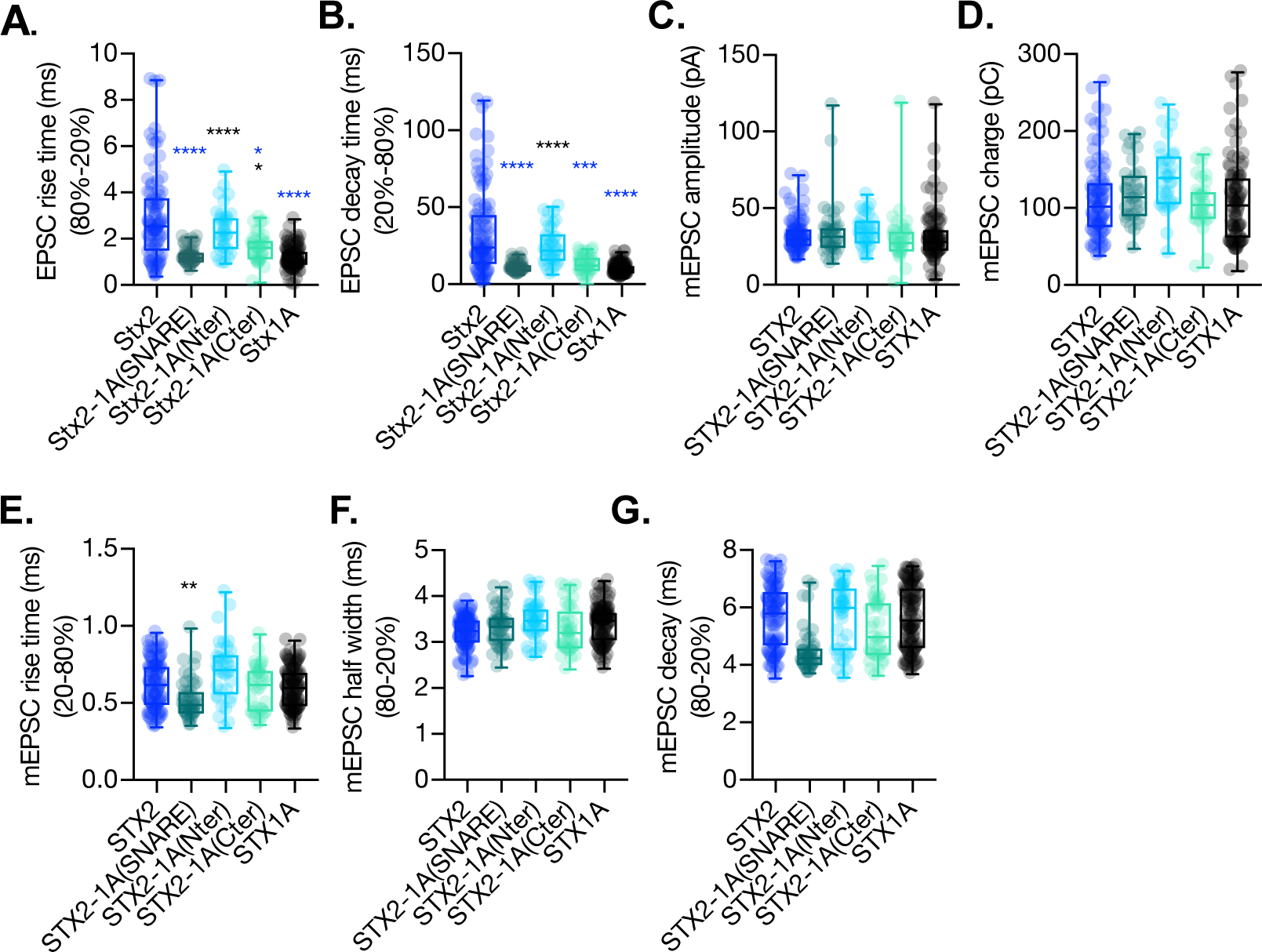
Quantification of kinetic parameters of the EPSC and the mEPSC of autaptic STX1A-null hippocampal mouse neurons rescued with STX2, STX2-1A(SNARE), STX2-1A(Nter) or STX2-1A(Cter). A. Quantification of the rise time (20-80%) of the EPSC. B. Quantification of the decay time (80-20%) of the EPSC. C. Quantification of the mEPSC charge. D. Quantification of the mEPSC amplitude. E. Quantification of the rise time (20-80%) of the mEPSC. F. Quantification of the half width of the mEPSC. G. Quantification of the decay time of the mEPSC. Data information: Data in (A-G) is show in a whisker-box plot. Each data point represents single observations, middle line represents the median, boxes represent the distribution of the data, where the majority of the data points lie and external data points represent outliers. Significances and P values of data were determined by non-parametric Kruskal-Wallis test followed by Dunn’s post hoc test or by Ordinary one-way ANOVA; **p≤0.05, **p≤0.01, ***p≤0.001, ****p≤0.0001. All data values are summarized in Figure 3 – Figure Supplement 1 – Source Data 1.

**Figure 6 – figure supplement 1.**
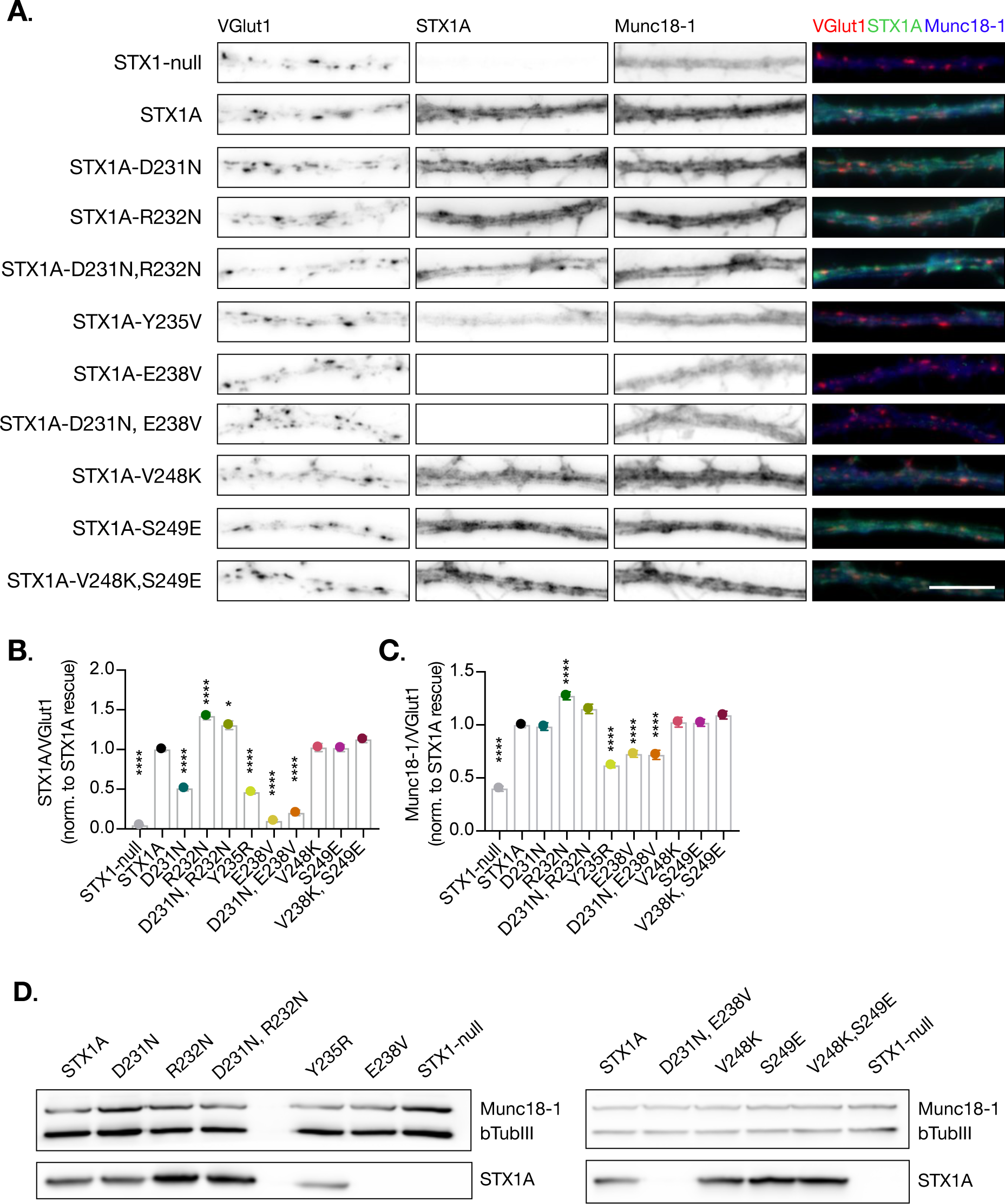
Quantification of STX2 levels at the synapse. A. Example images of Stx1-null neurons plated in high-density cultures and rescued with STX1A, STX2, STX2-1A(SNARE), STX2-1A(Nter), STX2-1A(Cter) or GFP (STX1-null) as negative control. Between DIV14-16 neurons were fixed. Neurons were stained with fluorophore-labeled antibodies that recognize VGlut1 (red in merge), STX1A (green in merge) and Munc18-1 (blue in merge) from left to right. Scale bar: 10μm. B. Quantification of the immunofluorescent intensity of STX1A normalized to the intensity of the same VGlut1-labeled ROIs. Values where normalized to STX1A WT values. C. Quantification of the immunofluorescent intensity of Munc18-1 normalized to the intensity of the same VGlut1-labeled ROIs. Values where normalized to STX1A WT values. D. SDS-PAGE of the electrophoretic analysis of neuronal lysates obtained from each experimental group. Proteins were detected using antibodies that recognize β-Tubuline III as loading control and STX1A and Munc18-1. Data information: In (B, C) data points represent the mean ±SEM. 7-9 images per group per culture were obtained and 3-5 ROI per image where analyzed. 3 replicates per group. Significances and P values of data were determined by non-parametric Kruskal-Wallis test followed by Dunn’s post hoc test; *p≤0.05, **p≤0.01, ***p≤0.001, ****p≤0.0001. All data values are summarized in Figure 6 – Figure Supplement 1 – Source Data 1.

## References

Abonyo, B. O., Gou, D., Wang, P., Narasaraju, T., Wang, Z., & Liu, L. (2004). Syntaxin 2 and SNAP-23 Are Required for Regulated Surfactant Secretion. Biochemistry, 43(12), 3499–3506. 10.1021/bi036338y

Arancillo, M., Min, S. W., Gerber, S., Münster-Wandowski, A., Wu, Y. J., Herman, M.,…Rosenmund, C. (2013). Titration of Syntaxin1 in mammalian synapses reveals multiple roles in vesicle docking, priming, and release probability. Journal of Neuroscience, 33(42), 16698–16714. 10.1523/JNEUROSCI.0187-13.2013

Bajohrs, M., Darios, F., Peak-Chew, S. Y., & Davletov, B. (2005). Promiscuous interaction of SNAP-25 with all plasma membrane syntaxins in a neuroendocrine cell. Biochemical Journal, 392(2), 283–289. 10.1042/BJ20050583

Baker, R. W., Jeffrey, P. D., Zick, M., Phillips, B. P., Wickner, W. T., & Hughson, F. M. (2015). A direct role for the Sec1/Munc18-family protein Vps33 as a template for SNARE assembly. Science, 349(6252), 1111–1114. 10.1126/science.aac7906

Basu, J., Betz, A., Brose, N., & Rosenmund, C. (2007). Munc13-1 C1 domain activation lowers the energy barrier for synaptic vesicle fusion. Journal of Neuroscience, 27(5), 1200–1210. 10.1523/JNEUROSCI.4908-06.2007

Bouazza-Arostegui, B., Camacho, M., Brockmann, M. M., Zobel, S., & Rosenmund, C. (2022). Deconstructing Synaptotagmin-1’s Distinct Roles in Synaptic Vesicle Priming and Neurotransmitter Release. Journal of Neuroscience, 42(14), 2866–2871. 10.1523/JNEUROSCI.1945-21.2022

Brunger, A. T. (2005). Structure and function of SNARE and SNARE-interacting proteins. Quarterly Reviews of Biophysics, 38(1), 1–47. 10.1017/S0033583505004051

Burkhardt, P., Hattendorf, D. A., Weis, W. I., & Fasshauer, D. (2008). Munc18a controls SNARE assembly through its interaction with the syntaxin N-peptide. EMBO Journal, 27(7), 923–933. 10.1038/emboj.2008.37

Chang, S., Trimbuch, T., & Rosenmund, C. (2018). Synaptotagmin-1 drives synchronous Ca 2+-triggered fusion by C 2 B-domain-mediated synaptic-vesicle-membrane attachment. Nature Neuroscience, 21(1), 33–42. 10.1038/s41593-017-0037-5

Chen, D., Minger, S. L., Honer, W. G., & Whiteheart, S. W. (1999). Organization of the secretory machinery in the rodent brain: Distribution of the t-SNAREs, SNAP-25 and SNAP-23. Brain Research, 831(1–2), 11–24. 10.1016/S0006-8993(99)01371-2

Dhara, M., Yarzagaray, A., Makke, M., Schindeldecker, B., Schwarz, Y., Shaaban, A.,…Bruns, D. (2016). v-SNARE transmembrane domains function as catalysts for vesicle fusion. ELife, 5, 1–25. 10.7554/elife.17571

Dolai, S., Liang, T., Orabi, A. I., Holmyard, D., Xie, L., Greitzer-Antes, D.,…Gaisano, H. Y. (2018). Pancreatitis-Induced Depletion of Syntaxin 2 Promotes Autophagy and Increases Basolateral Exocytosis. Gastroenterology, 154(6), 1805–1821.e5. 10.1053/j.gastro.2018.01.025

Dulubova, I., Khvotchev, M., Liu, S., Huryeva, I., Südhof, T. C., & Rizo, J. (2007). Munc18-1 binds directly to the neuronal SNARE complex. Proceedings of the National Academy of Sciences of the United States of America, 104(8), 2697–2702. 10.1073/pnas.0611318104

Fasshauer, D., Sutton, R. B., Brunger, A. T., & Jahn, R. (1998). Conserved structural features of the synaptic fusion complex: SNARE proteins reclassified as Q-and R-SNAREs. Proceedings of the National Academy of Sciences of the United States of America, 95(26), 15781–15786. 10.1073/pnas.95.26.15781

Fernandez, I., Ubach, J., Dulubova, I., Zhang, X., Südhof, T. C., & Rizo, J. (1998). Three-dimensional structure of an evolutionarily conserved N-terminal domain of syntaxin 1A. Cell, 94(6), 841–849. 10.1016/S0092-8674(00)81742-0

Gao, Y., Zorman, S., Gundersen, G., Xi, Z., Ma, L., Sirinakis, G.,…Zhang, Y. (2012). Single REconstituted Neuronal SNARE Complexes Zipper in Three Distinct Stages. Science, (September), 1340–1344.

Gerber, S. H., Rah, J. C., Min, S. W., Liu, X., De Wit, H., Dulubova, I.,…Südhof, T. C. (2008). Conformational switch of syntaxin-1 controls synaptic vesicle fusion. Science, 321(5895), 1507–1510. 10.1126/science.1163174

Hao, T., Feng, N., Gong, F., Yu, Y., Liu, J., & Ren, Y. X. (2023). Complexin-1 regulated assembly of single neuronal SNARE complex revealed by single-molecule optical tweezers. Communications Biology, 6(1), 1–12. 10.1038/s42003-023-04506-w

Hutt, D. M., Baltz, J. M., & Ngsee, J. K. (2005). Synaptotagmin VI and VIII and syntaxin 2 are essential for the mouse sperm acrosome reaction. Journal of Biological Chemistry, 280(21), 20197–20203. 10.1074/jbc.M412920200

Jiao, J., He, M., Port, S. A., Baker, R. W., Xu, Y., Qu, H.,…Zhang, Y. (2018). Munc18-1 catalyzes neuronal SNARE assembly by templating SNARE association. ELife, 7, 1–32. 10.7554/eLife.41771

Kádková, A., Murach, J., Pedersen, M. Ø., Malsam, A., Malsam, J., Söllner, T. H., & Sørensen, J. B. (2023). SNAP25 disease mutations change the energy landscape for synaptic exocytosis due to aberrant SNARE interactions. 10.1101/2023.05.21.541607

Khvotchev, M., Dulubova, I., Sun, J., Dai, H., Rizo, J., & Südhof, T. C. (2007). Dual modes of Munc18-1/SNARE interactions are coupled by functionally critical binding to syntaxin-1 N terminus. Journal of Neuroscience, 27(45), 12147–12155. 10.1523/JNEUROSCI.3655-07.2007

Lai, Y., Choi, U. B., Leitz, J., Rhee, H. J., Lee, C., Altas, B.,…Brunger, A. T. (2017). Molecular Mechanisms of Synaptic Vesicle Priming by Munc13 and Munc18. Neuron, 95(3), 591–607.e10. 10.1016/j.neuron.2017.07.004

Lois, C., Hong, E. J., Pease, S., Brown, E. J., & Baltimore, D. (2002). Germline transmission and tissue-specific expression of transgenes delivered by lentiviral vectors. Science, 295(5556), 868–872. 10.1126/science.1067081

Ma, C., Li, W., Xu, Y., & Rizo, J. (2011). Munc13 mediates the transition from the closed syntaxin-Munc18 complex to the SNARE complex. Nature Structural and Molecular Biology, 18(5), 542–549. 10.1038/nsmb.2047

Ma, L., Rebane, A. A., Yang, G., Xi, Z., Kang, Y., Gao, Y., & Zhang, Y. (2015). Munc18-1-regulated stage-wise SNARE assembly underlying synaptic exocytosis. ELife, 4, 1–31. 10.7554/elife.09580

McNew, J. A., Weber, T., Engelman, D. M., Söllner, T. H., & Rothman, J. E. (1999). The length of the flexible SNAREpin juxtamembrane region is a critical determinant of SNARE-dependent fusion. Molecular Cell, 4(3), 415–421. 10.1016/S1097-2765(00)80343-3

Meijer, M., Dörr, B., Lammertse, H. C., Blithikioti, C., Weering, J. R., Toonen, R. F.,…Verhage, M. (2018). Tyrosine phosphorylation of Munc18-1 inhibits synaptic transmission by preventing SNARE assembly. The EMBO Journal, 37(2), 300–320. 10.15252/embj.201796484

Oh, E., Kalwat, M. A., Kim, M. J., Verhage, M., & Thurmond, D. C. (2012). Munc18-1 regulates first-phase insulin release by promoting granule docking to multiple syntaxin isoforms. Journal of Biological Chemistry, 287(31), 25821–25833. 10.1074/jbc.M112.361501

Parisotto, D., Malsam, J., Scheutzow, A., Krause, J. M., & Söllner, T. H. (2012). SNAREpin assembly by Munc18-1 requires previous vesicle docking by synaptotagmin 1. Journal of Biological Chemistry, 287(37), 31041–31049. 10.1074/jbc.M112.386805

Peng, L., Liu, H., Ruan, H., Tepp, W. H., Stoothoff, W. H., Brown, R. H.,…Dong, M. (2013). Cytotoxicity of botulinum neurotoxins reveals a direct role of syntaxin 1 and SNAP-25 in neuron survival. Nature Communications, 4. 10.1038/ncomms2462

Quiñones, B., Riento, K., Olkkonen, V. M., Hardy, S., & Bennett, M. K. (1999). Syntaxin 2 splice variants exhibit differential expression patterns, biochemical properties and subcellular localizations. Journal of Cell Science, 112(23), 4291–4304. 10.1242/jcs.112.23.4291

RB, S., D, F., R, J., & AT, B. (1998). Crystal structure of a SNARE complex involved in synaptic exocytosis at A resolution. Nature, 395(September), 347–353.

Rizo, J. (2022). Molecular Mechanisms Underlying Neurotransmitter Release. Annual Review of Biophysics, 51, 377–408. 10.1146/annurev-biophys-111821-104732

Rizo, J., & Rosenmund, C. (2008). Synaptic vesicle fusion. Nature Structural and Molecular Biology, 15(7), 665–674. 10.1038/nsmb.1450

Rosenmund, C., & Stevens, C. F. (1996). Definition of the readily releasable pool of vesicles at hippocampal synapses. Neuron, 16(6), 1197–1207. 10.1016/S0896-6273(00)80146-4

Ruiter, M., Kádková, A., Scheutzow, A., Malsam, J., Söllner, T. H., & Sørensen, J. B. (2019). An Electrostatic Energy Barrier for SNARE-Dependent Spontaneous and Evoked Synaptic Transmission. Cell Reports, 26(9), 2340–2352.e5. 10.1016/j.celrep.2019.01.103

Schotten, S., Meijer, M., Walter, A. M., Huson, V., Mamer, L., Kalogreades, L.,…Cornelisse, L. N. (2015). Additive effects on the energy barrier for synaptic vesicle fusion cause supralinear effects on the vesicle fusion rate. ELife, 2015(4), 1–25. 10.7554/eLife.05531

Schupp, M., Malsam, J., Ruiter, M., Scheutzow, A., Wierda, K. D. B., Söllner, T. H., & Sørensen, J. B. (2016). Interactions between SNAP-25 and synaptotagmin-1 are involved in vesicle priming, clamping spontaneous and stimulating evoked neurotransmission. Journal of Neuroscience, 36(47), 11834–11836. 10.1523/JNEUROSCI.1011-16.2016

Shao, X., Li, C., Fernandez, I., Zhang, X., Südhof, T. C., & Rizo, J. (1997). Synaptotagmin–Syntaxin Interaction: The C2 Domain as a Ca2+-Dependent Electrostatic Switch. Neuron, 18(19), 5854–5860.

Sherry, D. M., Mitchell, R., Standifer, K. M., & du Plessis, B. (2006). Distribution of plasma membrane-associated syntaxins 1 through 4 indicates distinct trafficking functions in the synaptic layers of the mouse retina. BMC Neuroscience, 7, 1–25. 10.1186/1471-2202-7-54

Sørensen, J. B. (2009). Conflicting views on the membrane fusion machinery and the fusion pore. Annual Review of Cell and Developmental Biology, 25, 513–537. 10.1146/annurev.cellbio.24.110707.175239

Sørensen, J. B., Wiederhold, K., Müller, E. M., Milosevic, I., Nagy, G., De Groot, B. L.,…Fasshauer, D. (2006). Sequential N-To C-terminal SNARE complex assembly drives priming and fusion of secretory vesicles. EMBO Journal, 25(5), 955–966. 10.1038/sj.emboj.7601003

Stepien, K. P., Xu, J., Zhang, X., Bai, X. C., & Rizo, J. (2022). SNARE assembly enlightened by cryo-EM structures of a synaptobrevin–Munc18-1–syntaxin-1 complex. Science Advances, 8(25), 1–16. 10.1126/sciadv.abo5272

Südhof, T. C. (2013). Neurotransmitter release: The last millisecond in the life of a synaptic vesicle. Neuron, 80(3), 675–690. 10.1016/j.neuron.2013.10.022

Toonen, R. F. G., Wierda, K., Sons, M. S., De Wit, H., Cornelisse, L. N., Brussaard, A.,…Verhage, M. (2006). Munc18-1 expression level control synapse recovery by regulating readily releasable pool size. Proceedings of the National Academy of Sciences of the United States of America, 103(48), 18332–18337. 10.1073/pnas.0608507103

Trimbuch, T., Xu, J., Flaherty, D., Tomchick, D. R., Rizo, J., & Rosenmund, C. (2014). Re-examining how complexin inhibits neurotransmitter release: SNARE complex insertion or electrostatic hindrance? ELife, 2014(3), 1–28. 10.7554/eLife.02391

Vardar, G., Chang, S., Arancillo, M., Wu, Y. J., Trimbuch, T., & Rosenmund, C. (2016). Distinct functions of syntaxin-1 in neuronal maintenance, synaptic vesicle docking, and fusion in mouse neurons. Journal of Neuroscience, 36(30), 7911–7924. 10.1523/JNEUROSCI.1314-16.2016

Vardar, G., Salazar-Lázaro, A., Brockmann, M., Weber-Boyvat, M., Zobel, S., Kumbol, V. W. A.,…Rosenmund, C. (2021). Reexamination of n-terminal domains of syntaxin-1 in vesicle fusion from central murine synapses: Running title: Syntaxin-1 n-terminus in vesicle fusion. ELife, 10. 10.7554/eLife.69498

Vardar, G., Salazar-Lázaro, A., Zobel, S., Trimbuch, T., & Rosenmund, C. (2022). Syntaxin-1A modulates vesicle fusion in mammalian neurons via juxtamembrane domain dependent palmitoylation of its transmembrane domain. ELife, 11. 10.7554/eLife.78182

Verhage, M., Maia, A. S., Plomp, J. J., Brussaard, A. B., Heeroma, J. H., Vermeer, H.,…Südhof, T. C. (2000). Synaptic assembly of the brain in the absence of neurotransmitter secretion. Science, 287(5454), 864–869. 10.1126/science.287.5454.864

Voets, T., Toonen, R. F., Brian, E. C., De Wit, H., Moser, T., Rettig, J.,…Verhage, M. (2001). Munc18-1 promotes large dense-core vesicle docking. Neuron, 31(4), 581–592. 10.1016/S0896-6273(01)00391-9

Voleti, R., Jaczynska, K., & Rizo, J. (2020). Ca2+-dependent release of synaptotagmin-1 from the snare complex on phosphatidylinositol 4,5-bisphosphate-containing membranes. ELife, 9, 1–95. 10.7554/ELIFE.57154

Walter, A. M., Wiederhold, K., Bruns, D., Fasshauer, D., & Sørensen, J. B. (2010). Synaptobrevin N-terminally bound to syntaxin-SNAP-25 defines the primed vesicle state in regulated exocytosis. Journal of Cell Biology, 188(3), 401–413. 10.1083/jcb.200907018

Wang, S., Li, Y., Gong, J., Ye, S., Yang, X., Zhang, R., & Ma, C. (2019). Munc18 and Munc13 serve as a functional template to orchestrate neuronal SNARE complex assembly. Nature Communications, 10(1). 10.1038/s41467-018-08028-6

Wang, X., Gong, J., Zhu, L., Wang, S., Yang, X., Xu, Y.,…Ma, C. (2020). Munc13 activates the Munc18-1/syntaxin-1 complex and enables Munc18-1 to prime SNARE assembly. The EMBO Journal, 39(16), 1–22. 10.15252/embj.2019103631

Weber, J. P., Reim, K., & Sørensen, J. B. (2010). Opposing functions of two sub-domains of the SNARE-complex in neurotransmission. EMBO Journal, 29(15), 2477–2490. 10.1038/emboj.2010.130

Williams, D., Vicogne, J., Zaitseva, I., McLaughlin, S., & Perssin, J. E. (2009). Evidence that Electrostatic Interactions between Vesicle-associated Membrane Protein 2 and Acidic Phospholipids May Modulate the Fusion of Transport Vesicles with the Plasma Membrane. Molecular Biology of the Cell, 20, 4524–4530. 10.1091/mbc.E09

Wu, Y. J., Tejero, R., Arancillo, M., Vardar, G., Korotkova, T., Kintscher, M.,…Rosenmund, C. (2015). Syntaxin 1B is important for mouse postnatal survival and proper synaptic function at the mouse neuromuscular junctions. Journal of Neurophysiology, 114(4), 2404–2417. 10.1152/jn.00577.2015

Xu, J., Mashimo, T., & Südhof, T. C. (2007). Synaptotagmin-1,-2, and −9: Ca2+ Sensors for Fast Release that Specify Distinct Presynaptic Properties in Subsets of Neurons. Neuron, 54(4), 567–581. 10.1016/j.neuron.2007.05.004

Xue, M., Lin, Y. Q., Pan, H., Reim, K., Deng, H., Bellen, H. J., & Rosenmund, C. (2009). Tilting the Balance between Facilitatory and Inhibitory Functions of Mammalian and Drosophila Complexins Orchestrates Synaptic Vesicle Exocytosis. Neuron, 64(3), 367–380. 10.1016/j.neuron.2009.09.043

Xue, M., Reim, K., Chen, X., Chao, H. T., Deng, H., Rizo, J.,…Rosenmund, C. (2007). Distinct domains of complexin I differentially regulate neurotransmitter release. Nature Structural and Molecular Biology, 14(10), 949–958. 10.1038/nsmb1292

Zhang, Y. (2017). Energetics, kinetics, and pathway of SNARE folding and assembly revealed by optical tweezers. Protein Science, 26(7), 1252–1265. 10.1002/pro.3116

Zhang, Y., Ma, L., & Bao, H. (2022). Energetics, kinetics, and pathways of SNARE assembly in membrane fusion. Critical Reviews in Biochemistry and Molecular Biology, 57(4), 443–460. 10.1080/10409238.2022.2121804

Zhou, P., Pang, Z. P., Yang, X., Zhang, Y., Rosenmund, C., Bacaj, T., & Südhof, T. C. (2013). Syntaxin-1 N-peptide and H abc-domain perform distinct essential functions in synaptic vesicle fusion. EMBO Journal, 32(1), 159–171. 10.1038/emboj.2012.307

Zhou, Q., Lai, Y., Bacaj, T., Zhao, M., Lyubimov, A. Y., Uervirojnangkoorn, M.,…Brunger, A. T. (2015). Architecture of the synaptotagmin-SNARE machinery for neuronal exocytosis. Nature, 525(7567), 62–67. 10.1038/nature14975

Zhou, Q., Zhou, P., Wang, A. L., Wu, D., Zhao, M., Südhof, T. C., & Brunger, A. T. (2017). The primed SNARE-complexin-synaptotagmin complex for neuronal exocytosis. Nature, 548(7668), 420–425. 10.1038/nature23484

